# ENPP1 is an innate immune checkpoint of the anticancer cGAMP-STING pathway

**DOI:** 10.1101/2023.06.01.543353

**Authors:** Songnan Wang, Volker Böhnert, Alby J. Joseph, Valentino Sudaryo, Jason Swinderman, Feiqiao B.Yu, Xuchao Lyu, Gemini Skariah, Vishvak Subramanyam, Luke A. Gilbert, Hani Goodarzi, Lingyin Li

## Abstract

ENPP1 expression correlates with poor prognosis in many cancers, and we previously discovered that ENPP1 is the dominant hydrolase of extracellular cGAMP: a cancer-cell-produced immunotransmitter that activates the anticancer STING pathway. However, ENPP1 has other catalytic activities and the molecular and cellular mechanisms contributing to its tumorigenic effects remain unclear. Here, using single cell RNA-seq (scRNA-seq), we show that ENPP1 overexpression drives primary breast tumor growth and metastasis by synergistically dampening extracellular cGAMP-STING mediated antitumoral immunity and activating immunosuppressive extracellular adenosine (eADO) signaling. In addition to cancer cells, stromal and immune cells in the tumor microenvironment (TME) also express ENPP1 that restrains their response to tumor-derived cGAMP. *Enpp1* loss-of-function in both cancer cells and normal tissues slowed primary tumor initiation and growth and prevented metastasis in an extracellular cGAMP- and STING-dependent manner. Selectively abolishing the cGAMP hydrolysis activity of ENPP1 phenocopied total ENPP1 knockout, demonstrating that restoration of paracrine cGAMP-STING signaling is the dominant anti-cancer mechanism of ENPP1 inhibition. Strikingly, we find that breast cancer patients with low *ENPP1* expression have significantly higher immune infiltration and improved response to therapeutics impacting cancer immunity upstream or downstream of the cGAMP-STING pathway, like PARP inhibitors and anti-PD1. Altogether, selective inhibition of ENPP1’s cGAMP hydrolase activity alleviates an innate immune checkpoint to boost cancer immunity and is therefore a promising therapeutic approach against breast cancer that may synergize with other cancer immunotherapies.

## Introduction

The strategy of blocking adaptive immune checkpoints (PD-1, PD-L1, and/or CTLA-4) offers curative immunotherapy for some patients with otherwise terminal cancer diagnoses – however, only a minority of patients respond to immune checkpoint blockade (ICB) therapy and many cancer types remain inaccessible by this treatment. (Hodi *et al*., 2010; Brahmer *et al*., 2015; Garon *et al*., 2015; Ribas and Wolchok, 2018; Haslam and Prasad, 2019; Schmid *et al*., 2020). ICB resistance can occur through a variety of mechanisms, one of which is insufficient lymphocyte infiltration into the tumor, which is determined by our innate immune system’s ability to detect and communicate the presence of cancer (Fuertes *et al*., 2011; Woo, Corrales and Gajewski, 2015; Corrales *et al*., 2017). Cancer cells have a variety of strategies for suppressing innate immunity (Willingham *et al*., 2012; André *et al*., 2018; Barkal *et al*., 2018; Zhang *et al*., 2018); therefore, a deep understanding of innate immune checkpoints holds the potential to unlock the full power of immunotherapy against immunologically “cold” tumors.

The cytosolic double-stranded DNA (dsDNA)-sensing STING pathway is a key innate immune pathway that detects and responds to chromosomal instability (CIN) and extrachromosomal DNA present in cancer cells (Harding *et al*., 2017; MacKenzie *et al*., 2017; Turner *et al*., 2017; Bakhoum *et al*., 2018a). Cytosolic dsDNA is detected by the enzyme cyclic-GMP-AMP synthase (cGAS) (Sun *et al*., 2013), which synthesizes the cyclic dinucleotide cGAMP (Ablasser *et al*., 2013; Wu *et al*., 2013). cGAMP binds and activates STING, leading to production of type I interferons (IFN-I) and downstream immune cell infiltration (Corrales *et al*., 2015; Wang *et al*., 2017). We and others discovered that cancer cells produce and secrete cGAMP into the extracellular space (Carozza, Böhnert, *et al*., 2020; Maltbaek, Snyder and Stetson, 2021), which is then taken up by surrounding host cells, leading to paracrine STING activation (Luteijn *et al*., 2019; Ritchie *et al*., 2019; Lahey, Wen, *et al*., 2020; Zhou *et al*., 2020; Cordova *et al*., 2021). We found that extracellular cGAMP is important for the curative effect of ionizing radiation (IR) in a murine breast cancer model (Carozza, Böhnert, *et al*., 2020). This serves as a hint of the importance of the extracellular cGAMP-STING axis in cancer. However, unlike cell-intrinsic cGAS-STING signaling, understanding of paracrine cGAS-STING signaling in cancer is only in its nascency.

One important negative regulator of the extracellular cGAMP-STING pathway is Ectonucleotide Pyrophosphatase/Phosphodiesterase (ENPP1), the dominant hydrolase that degrades extracellular cGAMP (Li *et al*., 2014; Carozza, Böhnert, *et al*., 2020). There is mounting evidence that ENPP1 promotes cancer in humans (Umar *et al*., 2009; Li *et al*., 2021), and our previous work showed that ENPP1 inhibitors have efficacy in murine models of breast and pancreatic cancers (Carozza, Böhnert, *et al*., 2020; Carozza, Brown, *et al*., 2020). Despite the phenotypic link, ENPP1’s dominant mechanisms of action in cancer remain unclear, particularly because ENPP1 may impact multiple aspects of cancer signaling and metabolism. In addition to degrading extracellular cGAMP, ENPP1 also produces immunosuppressive extracellular adenosine (eADO) as a downstream metabolite of cGAMP hydrolysis, which has been linked to metastasis of murine CIN-high breast cancers (Li *et al*., 2021) and self-seeding of circulating breast tumor cells (De Córdoba *et al*., 2022). Therefore, numerous questions remain surrounding the molecular and cellular mechanisms of ENPP1’s tumorigenic activity.

In this study, we dissected ENPP1’s molecular and cellular mechanisms and characterized its downstream impacts in breast cancer. We report for the first time a causal link between ENPP1’s roles in breast cancer onset, progression, and metastasis and its antagonism of the cGAMP-STING mediated antitumoral immunity. Our results conclusively show that inhibiting cGAMP hydrolysis activity of ENPP1 is necessary and sufficient to block its tumorigenic effect. Our study demonstrates the paramount importance of paracrine STING activation in protecting against primary tumor development and metastasis. Taken together, we conclude that ENPP1 is an innate immune checkpoint restraining STING pathway activation in the TME, and that therapeutically targeting ENPP1’s cGAMP hydrolysis activity could boost anti-tumor immunity.

## Results

### ENPP1’s catalytic activity drives breast tumor growth and metastasis by restricting adaptive immune infiltration

ENPP1 expression levels have been shown to correlate with poor prognosis in several cancer types. Patients in the METABRIC database with breast tumors expressing high levels of *ENPP1* mRNA have a significantly worse disease-free survival rate, despite exhibiting a similar distribution across disease stages as the ENPP1-low group (**Figure 1A**). Furthermore, patients with stage IV metastatic disease have significantly higher *ENPP1* RNA expression than patients with stage III disease (**Figure 1B**). To test if we can recapitulate the tumorigenic effects of ENPP1 in mice, we performed orthotopic implantation of *Enpp1*-knockout 4T1 murine breast cancer cells overexpressing either WT ENPP1 (ENPP1^WT-OE^) or catalytically dead ENPP1 (ENPP1^T238A-OE^) (Kato *et al*., 2018) into WT mice **(Figure S1A-C)**. ENPP1^WT-OE^ 4T1 cells exhibited faster primary tumor growth and more lung metastases (**Figure 1C, S1E-F**) without affecting cell proliferation (**Figure S1D**), implicating a non-tumor cell intrinsic mechanism of enhanced tumor growth and metastasis. To thoroughly characterize the impact of ENPP1 overexpression on the tumor microenvironment (TME), we performed single-cell RNA-seq (scRNA-seq) on primary tumors and lungs colonized by metastases collected from this experiment. After strict quality control and filtration, we collected a total of 32,539 cells. We performed unsupervised graph-based clustering on all cells and identified 18 major clusters based on established cell markers (**Figure 1D, S2A-B**).

**Figure 1.**
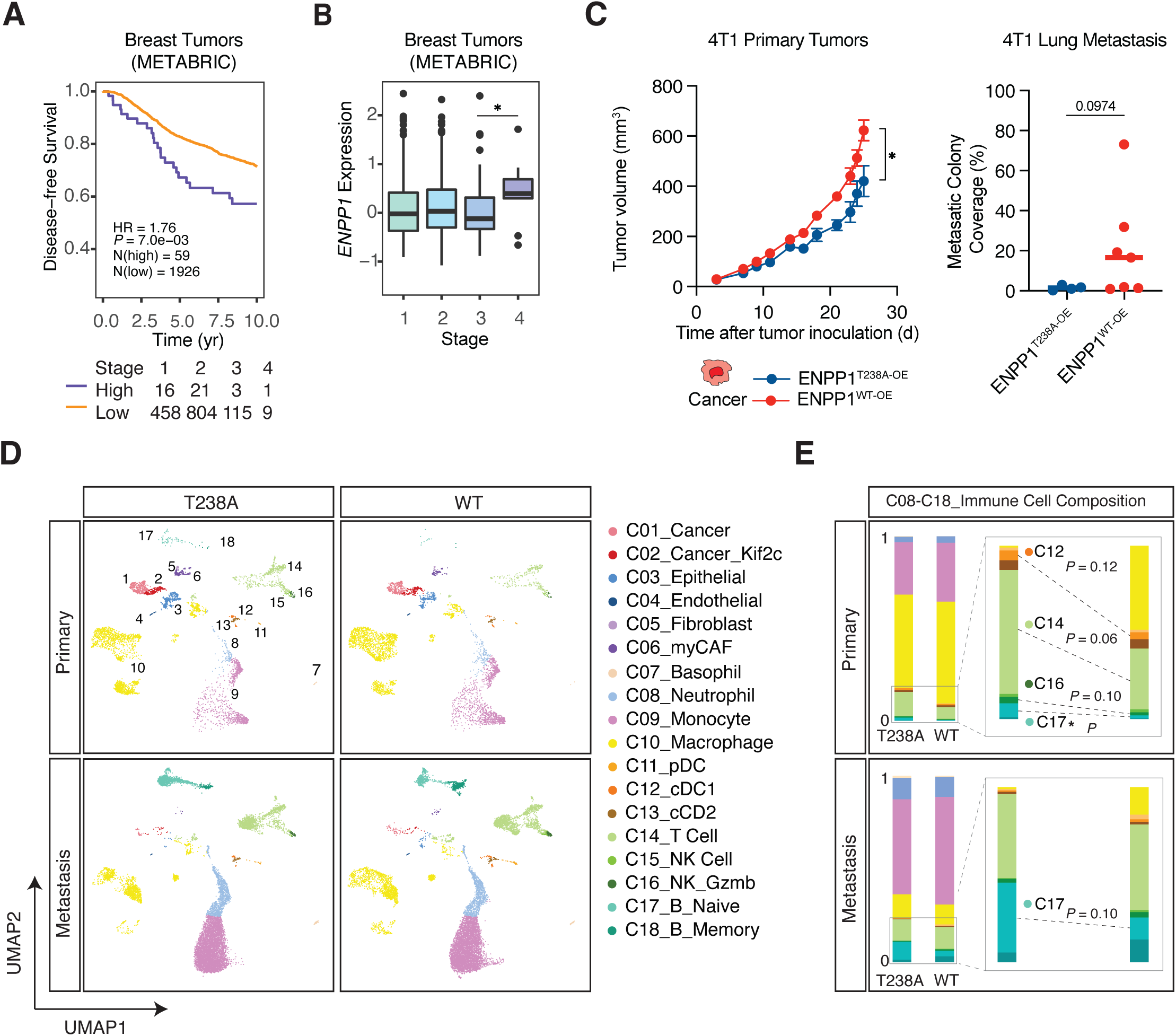
ENPP1’s catalytic activity drives breast tumor growth and metastasis by restricting immune infiltration. (A) Disease-free survival of breast cancer patients in the METABRIC database in the *ENPP1*-high group (n = 59) and *ENPP1*-low group (n = 1926) and the number of patients stratified by stages. Threshold for high versus low expression was set at which *P* value was the smallest. (B) *ENPP1* expression in patients with stage 1-4 breast cancer. Shown as box plots of median and interquartile levels. *P* value was determined by the nonparametric Mann-Whitney *U* test. (C) Primary tumor volumes and quantification of lung metastases of WT BALB/cJ mice bearing ENPP1^T238A-OE^ and ENPP1^WT-OE^ 4T1 tumors (n = 5 and 9 mice). Tumor growth curves were plotted as mean ± SEM. Metastasis data were plotted as mean. *P* values were determined by unpaired *t* test with Welch correction. (D) UMAP plots of the annotated clusters of ENPP1^T238A-OE^ and ENPP1^WT-OE^ 4T1 primary tumors and metastasis colonized lungs. (E) Barplots comparing immune cell compositions (containing C08-C18) between ENPP1^T238A-OE^ and ENPP1^WT-OE^ 4T1 primary tumors and metastasis colonized lungs. \**P* ≤ 0.05.; *P* value is shown if it is between 0.05 - 0.15. See also Figure S1 and S2.

Although the ENPP1^WT-OE^ condition did not alter the composition of non-immune cells (**Figure S2C**), it led to a decreased proportion of conventional DC type 1 (cDC1s), T cells, and *Gzmb*^+^ cytotoxic NK cells (**Figure 1E**). Unexpectedly, overexpression of WT ENPP1 resulted in a dramatic decrease in tumor infiltrating naïve B cells in both the primary and metastatic TME compared with the catalytic mutant (**Figure 1E**). Together, we conclude that ENPP1’s catalytic activity restricts adaptive immune cell infiltration, contributing to its role in promoting breast cancer growth and metastasis.

### ENPP1’s catalytic activity promotes immune suppression in primary tumors and lung metastases

Building on our findings that ENPP1 catalytic activity alters the composition of the tumor immune compartment, we next investigated its impact on the functional landscape of tumor-infiltrating immune cells. We first turned our attention to innate immune cells. We found increased expression of Arginase 1 (*Arg1*) in monocytes and macrophages in mice injected with ENPP1^WT-OE^ 4T1 cells (**Figure 2A, S2D**), which is associated with pro-tumor myeloid-derived suppressor cells (MDSCs) (Rodriguez *et al*., 2004) and M2-like macrophages (Yang and Ming, 2014), respectively. We further categorized macrophages into four subtypes based on expression of reported macrophage identity markers (Cheng *et al*., 2021) and functional markers (**Figure S3A, B**). These four tumor-associated macrophage (TAM) subsets express different levels of anti-tumor M1-like macrophage markers versus pro-tumor M2-like macrophage markers, with *Vcan*^+^ TAMs being the most M1-like, and *Pparg^+^* being exclusively immunosuppressive (**Figure S3B**). We observed a decrease in *Vcan*^+^ and *C1qc^+^* TAMs and a concomitant increase in M2-like *Spp1^+^* TAMs in ENPP1^WT-OE^, suggesting that ENPP1 catalytic activity favors the polarization of M1-like to M2-like TAMs (**Figure S3C**). This is consistent with our previously reported effect of extracellular cGAMP depletion on macrophage polarization (Cordova *et al*., 2021).

**Figure 2.**
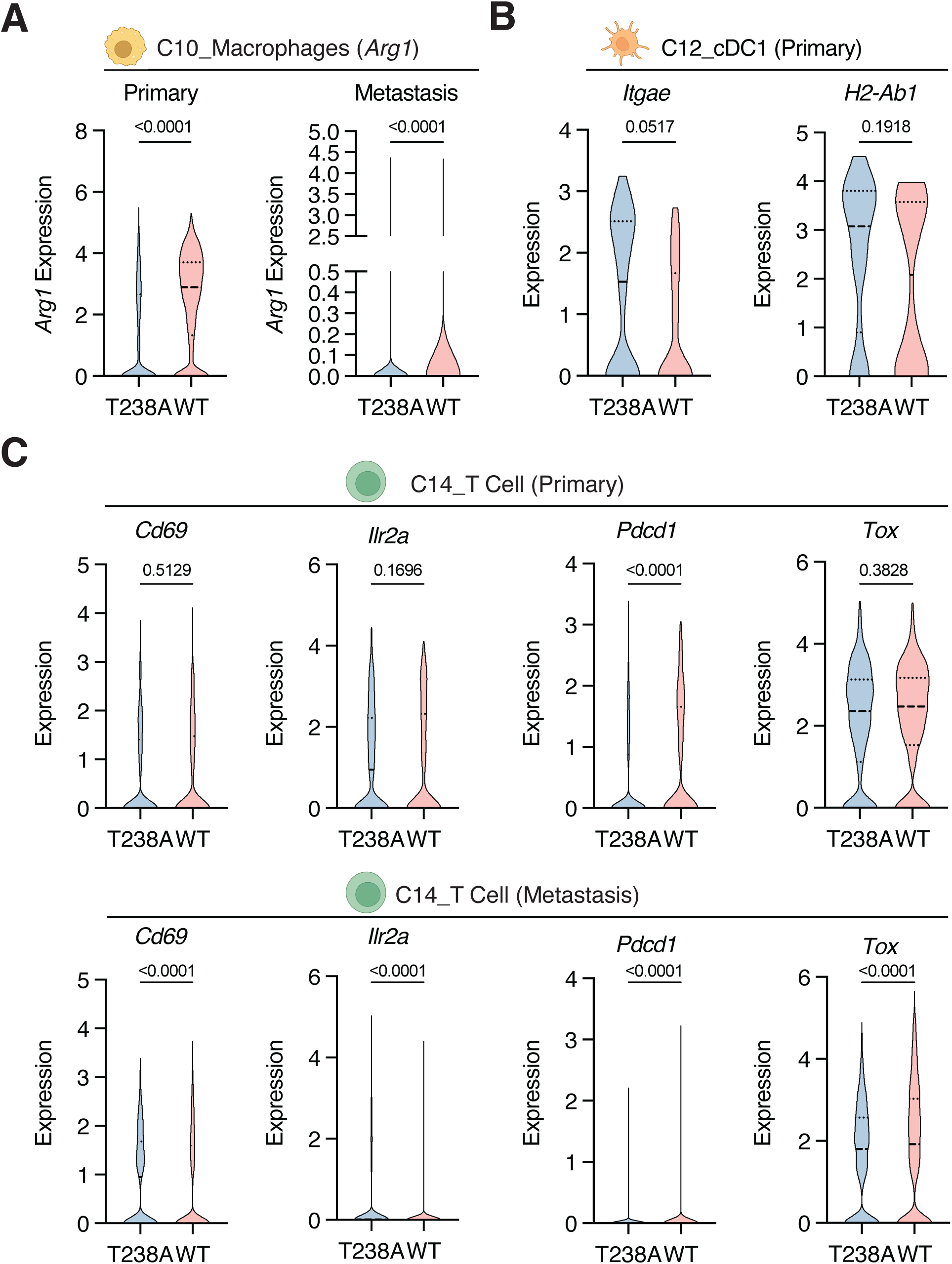
ENPP1’s catalytic activity promotes immune suppression in primary tumors and lung metastases. (A-C) Violin plots of indicated transcripts in indicated cell types comparing between ENPP1^T238A-OE^ and ENPP1^WT-OE^ 4T1 tumors or metastases. *Arg1* in macrophages (C10) in primary tumors and lung metastases (A); *Itgae* and *H2-Ab1* in cDC1s (C12) in primary tumors (B); *Cd69*, *Ilr2a*, *Pdcd1*, and *Tox* in T cells (C14) in primary tumors and lung metastases (C). *P* values were determined by nonparametric Mann-Whitney *U* test. cDC1 stands for conventional dendritic cell type 1. See also Figure S2 and S3

Focusing on antigen presenting cells (APCs) that orchestrate innate-adaptive crosstalk, we noticed a decrease in *Itgae* expressing migratory cDC1 in primary tumors, but not metastases, in the presence of ENPP1 catalytic activity (**Figure 2C, S2E**), mirroring the effects we previously observed with extracellular cGAMP depletion during ionizing radiation (IR) treatment (Carozza, Böhnert, *et al*., 2020). Additionally, cDC1s in ENPP1^WT-OE^ primary tumors, but not metastases, express less *H2-Ab1*, suggesting decreased antigen-presentation capacity (**Figure 2C, S2E**). We reason that APC recruitment and education at the initial site of cancer encounter is important in mounting a successful adaptive immune response, and this process appears to be diminished when tumors overexpress ENPP1.

Examining changes in adaptive immune cells, we found that the relative abundance of T cell subpopulations was not significantly altered between the WT and catalytic mutant ENPP1 conditions (**Figure S3D-F**). However, compared to ENPP1^T238A-OE^, WT ENPP1 activity decreased the expression of *Cd69* (an early T cell activation marker) and *Ilr2a* (a late T cell activation marker), and increased the expression of *Pdcd1* and *Tox* (exhausted T cell markers) in the T cells infiltrating both primary tumors and lung metastases (**Figure 2E**). In summary, our scRNA-seq data suggests that cancer cell-derived ENPP1 catalytic activity attenuated T cell activation while promoting exhaustion in primary tumors and sites of metastasis. Taken together, ENPP1 catalytic activity shapes the immunosuppressive TME both in primary tumors and metastases.

### ENPP1 inhibits cGAMP-STING signaling and increases eADO signaling to suppress anti-tumor immunity

To understand the mechanisms of cancer-derived ENPP1 overexpression in driving immunosuppression, we first analyzed expression of genes in the extracellular cGAMP-STING pathway (**Figure 3A-B, S4A**). Notably, *Cgas* expression is the highest in a distinct cancer cell cluster annotated for overexpressing *Kif2c*, which has been reported to drive CIN and metastasis (Bakhoum *et al*., 2018a). In contrast, *Tmem173/Sting* is expressed at relatively high levels in endothelial cells, fibroblasts, macrophages, and DCs (**Figure 3A**). The differential expression of cGAS and STING supports our model that cancers produce and secrete cGAMP, which is then detected by surrounding host cells (Carozza, Böhnert, *et al*., 2020). We defined potential cGAMP responder cells as those with low expression of *Cgas* and high expression of *Sting* and interferon-stimulated genes (ISGs) *Ifitm1*, *Ifitm2*, and *Ifitm3* (**Figure 3B, S4A**), which matched well with our previous report of cell types that respond to extracellular cGAMP (Cordova *et al*., 2021). We then examined expression of the only known murine cGAMP transporters, LRRC8A:C and LRRC8A:E (Lahey, Mardjuki, *et al*., 2020; Zhou *et al*., 2020), in these responder cells. While endothelial cells, macrophages, and DCs express genes encoding the LRRC8A:C complex, fibroblasts uniquely express genes encoding the LRRC8A:E complex (**Figure 3B**).

**Figure 3.**
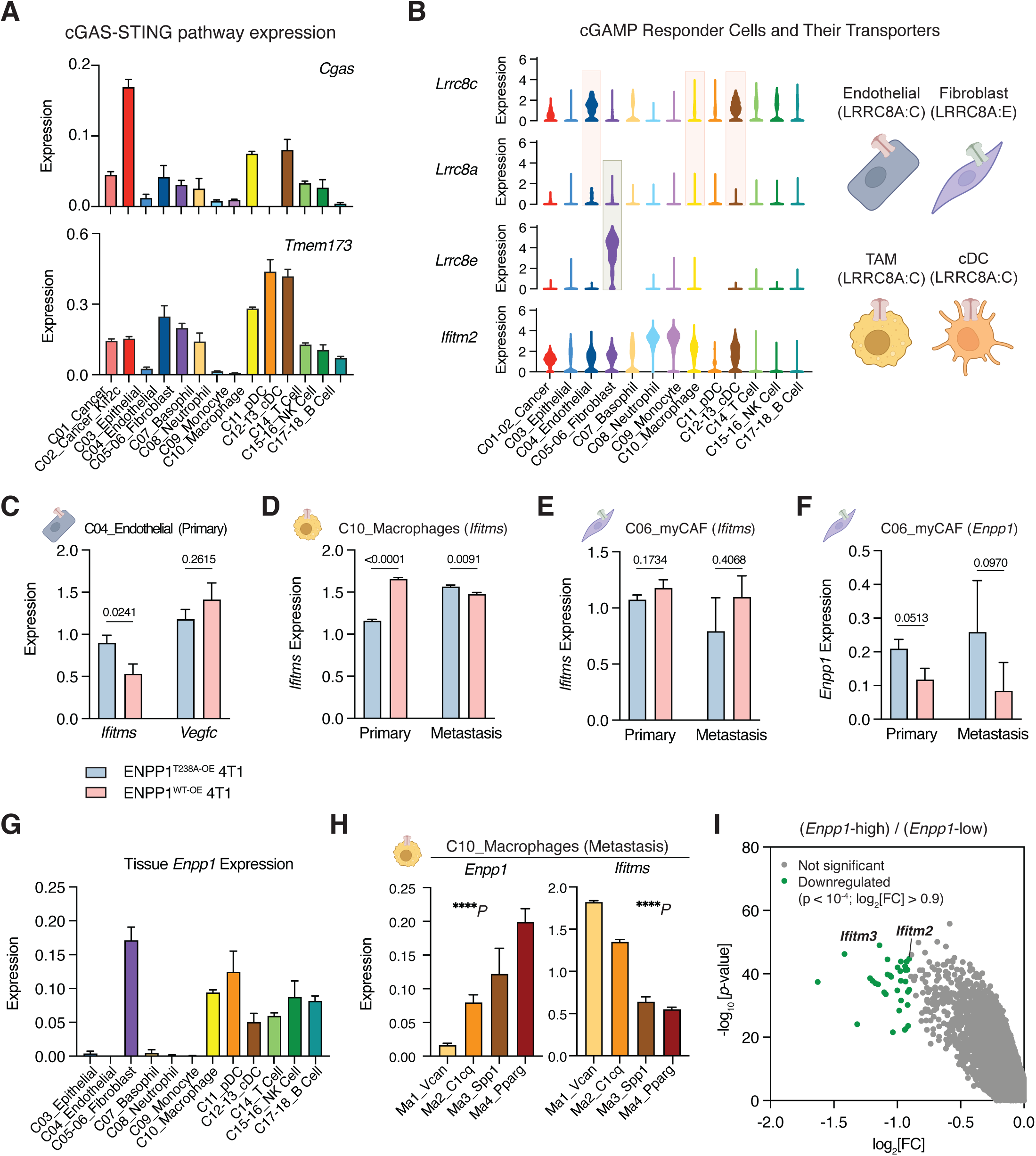
ENPP1 expressed on cancer and responder cells blocks paracrine cGAMP-STING activation. (A) Bar graphs of *Cgas* and *Tmem173* expression across the annotated clusters. (B) Violin plots of *Lrr8cc*, *Lrrc8ca*, *Lrrc8ce* and *Ifitm2* across the annotated clusters. Schematic of cGAMP responder cells and their putative transporters: LRRC8A:C in endothelial cells (C04), TAM (C10) and cDC (C11-12); LRRC8A:E in fibroblasts (C05-06). (C-F) Bar graphs of indicated transcripts in indicated cell types comparing between ENPP1^T238A-OE^ and ENPP1^WT-OE^ 4T1 tumors or metastases. *Ifitms* and *Vegfc* in endothelial cells (C04) in primary tumors (C); *Ifitms* in macrophages (C10) in primary tumors and metastases (D); *Ifitms* in myCAFs (C06) in primary tumors and metastases (E); *Enpp1* in myCAFs (C06) in primary tumors and metastases (F). *P* values in (C-D) were determined by nonparametric Mann-Whitney *U* test. (G) Bar graphs of *Enpp1* expression across the annotated clusters. (H) Bar graphs of *Enpp1* and *Ifitms* across the annotated macrophage subclusters. P values were determined by ordinary one-way ANOVA test. (I) Differentially expressed genes in *Enpp1*-high vs*. Enpp1*-low groups. Bars represent mean ± SEM. TAM stands for tumor-associated macrophages. cDC stands for conventional dendritic cell. myCAF stands for myofibroblastic cancer-associated fibroblast. See also Figure S4 and S5.

We next examined how overexpression of WT ENPP1 in cancer cells affected STING activation in cGAMP responder cells as measured by their combined *Ifitm1*, *Ifitm2*, and *Ifitm3* expression (*Ifitms*). Looking first at endothelial cells, we found that elevated ENPP1 catalytic activity from ENPP1^WT-OE^ 4T1s suppressed endothelial *Ifitms* expression, with a more pronounced effect in primary tumors than lung metastases (**Figure 3C, S4B**). Interferon (IFN) signaling is known to downregulate vascular endothelial growth factors (VEGFs) and inhibit angiogenesis (von Marschall *et al*., 2003); we observed that endothelial cells with blunted IFN signaling in ENPP1^WT-OE^ primary tumors express more *Vegfc* (**Figure 3C, S4B**). Turning our focus to TAMs, we show that in metastasis, ENPP1^WT-OE^ 4T1s with increased cGAMP degradation activity led to decreased expression of *Ifitms* as expected (**Figure 3D**). However, we observed the opposite trend in primary tumors where *Ifitms* expression in TAMs increased in ENPP1^WT-OE^ tumors (**Figure 3D**). We noticed that *Lrrc8c* expression in TAMs is also higher in ENPP1^WT-OE^ primary tumors but not metastases (**Figure S4C**). It is possible that increased cGAMP import activity in ENPP1^WT-OE^ primary tumors contributes to the unexpected increase in their *Ifitms* expression.

To understand how ENPP1^WT-OE^ 4T1s promote immunosuppressive phenotypes in primary tumor-resident TAMs despite increased ISG expression in these cells, we examined the activation status of the eADO pathway: a potential STING-independent downstream effect of cGAMP hydrolysis (**Figure S5A**). We observed increased expression of *Adora2b* (the eADO receptor), *Tgfb1* and *Il10rb* (downstream of eADO signaling), and *Hp* (a gene that is transcriptionally upregulated by adenosine signaling) in primary tumors (Boison and Yegutkin, 2019; Hatfield *et al*., 2019; De Córdoba *et al*., 2022), indicating activation of the eADO pathway specifically in TAMs in primary tumors but not in metastases (**Figure S5B**). Haptoglobin (HP) secretion by cancer cells upon of eADO signaling has been reported to recruit polymorphonuclear MDSCs (PMN-MDSCs) and promote self-seeding of ENPP1-high circulating tumor cells (CTCs) (De Córdoba *et al*., 2022). However, we did not observe increased *Hp* expression in ENPP1^WT-OE^ cancer cells (**Figure S5C**). Instead, neutrophils and monocytes in the metastatic niche had the biggest increase and the highest overall expression of *Hp* in ENPP1^WT-OE^ tumors (**Figure S5C**). Together, our data support a model in which ENPP1 overexpression promotes primary tumor growth and metastasis through synergistic stimulation of the eADO pathway and inhibition of the cGAMP-STING pathway (**Figure S5D**).

### ENPP1 expressed on host responder cells suppresses paracrine STING activation

We next turned our attention to a third class of cGAMP responder cells, myofibroblast-like cancer-associated fibroblasts (myCAFs): a subtype of fibroblasts with immunosuppressive functions (Mhaidly and Mechta-Grigoriou, 2020). Like TAMs, myCAFs also exhibited an unexpected decrease in STING pathway activation as measured by *Ifitms* expression, with a concomitant increase in *Enpp1* expression, in ENPP1^T238A-OE^ primary tumors (**Figure 3E,F**). Previous studies and our analysis so far have focused on ENPP1 expressed by cancer cells. However, these data hint at the importance of the *Enpp1* level expressed by a responder cell in regulating its paracrine STING signaling.

To build a more comprehensive picture of *Enpp1* expression across host cells in the TME, we examined its expression level and downstream impact on STING activation in stromal and immune cells. Among the responder cells, we found that fibroblasts, macrophages, and DCs express relatively high levels of *Enpp1* (**Figure 3G**). Endothelial cells, on the other hand, do not express measurable levels of *Enpp1* (**Figure 3G**), which could explain the predominant effect of cancer-derived ENPP1 on their STING activation profile (**Figure 3C**). Additionally, we found that *Ifitms* expression in subpopulations of TAMs in the metastases anti-correlates with their *Enpp1* expression level (**Figure 3H**). Specifically, there is a step-wise increase in *Enpp1* expression and a corresponding decrease in *Ifitms* expression from the immunostimulatory M1-like macrophages to the immunosuppressive M2-like macrophages (**Figure 3H**). This result could potentially explain our previous findings that M2-polarized macrophages are less sensitive to cancer-derived extracellular cGAMP than M1-polarized macrophages (Cordova *et al*., 2021).

Expanding this correlation to other cell types, we performed differential gene expression analysis between *Enpp1*-high cells (Enpp1 > 1.24, 338 cells) and *Enpp1*-low cells (0 < *Enpp1* < 1.24, 1040 cells) and found that *Ifitm2* and *Ifitm3* are among the most significantly downregulated genes in *Enpp1*-high cells (**Figure 3I**). Together, our results demonstrate that ENPP1 expressed on cGAMP responder cells potently inhibits these cells’ paracrine STING activation. These findings suggest the need to inhibit both cancer- and host-derived ENPP1 to alleviate its tumorigenic effect and emphasize the impact of host ENPP1 expression level in different cell types on the downstream effects of paracrine STING signaling.

### *Enpp1* knockout in cancer and tissue cells additively delay tumor growth and metastasis

Next, we formally tested the relative contribution of cancer- and host-derived ENPP1 on breast tumor growth and metastasis. We implanted WT or *Enpp1^-/-^* 4T1 cells into WT or *Enpp1^-/-^* BALB/cJ mice and treated established tumors (palpable around 100 mm^3^) with ionizing radiation (IR) to further induce cGAMP production (Carozza, Böhnert, *et al*., 2020). Indeed, depleting cancer- or tissue-derived ENPP1 had an additive effect on slowing tumor growth, with tissue ENPP1 playing a larger role (**Figure 4A**). Cancer- and host-derived ENPP1 contribute equally to intratumoral cGAMP degradation activity, while tissue-derived ENPP1 is mainly responsible for cGAMP degradation in serum (**Figure S6A-B**).

**Figure 4.**
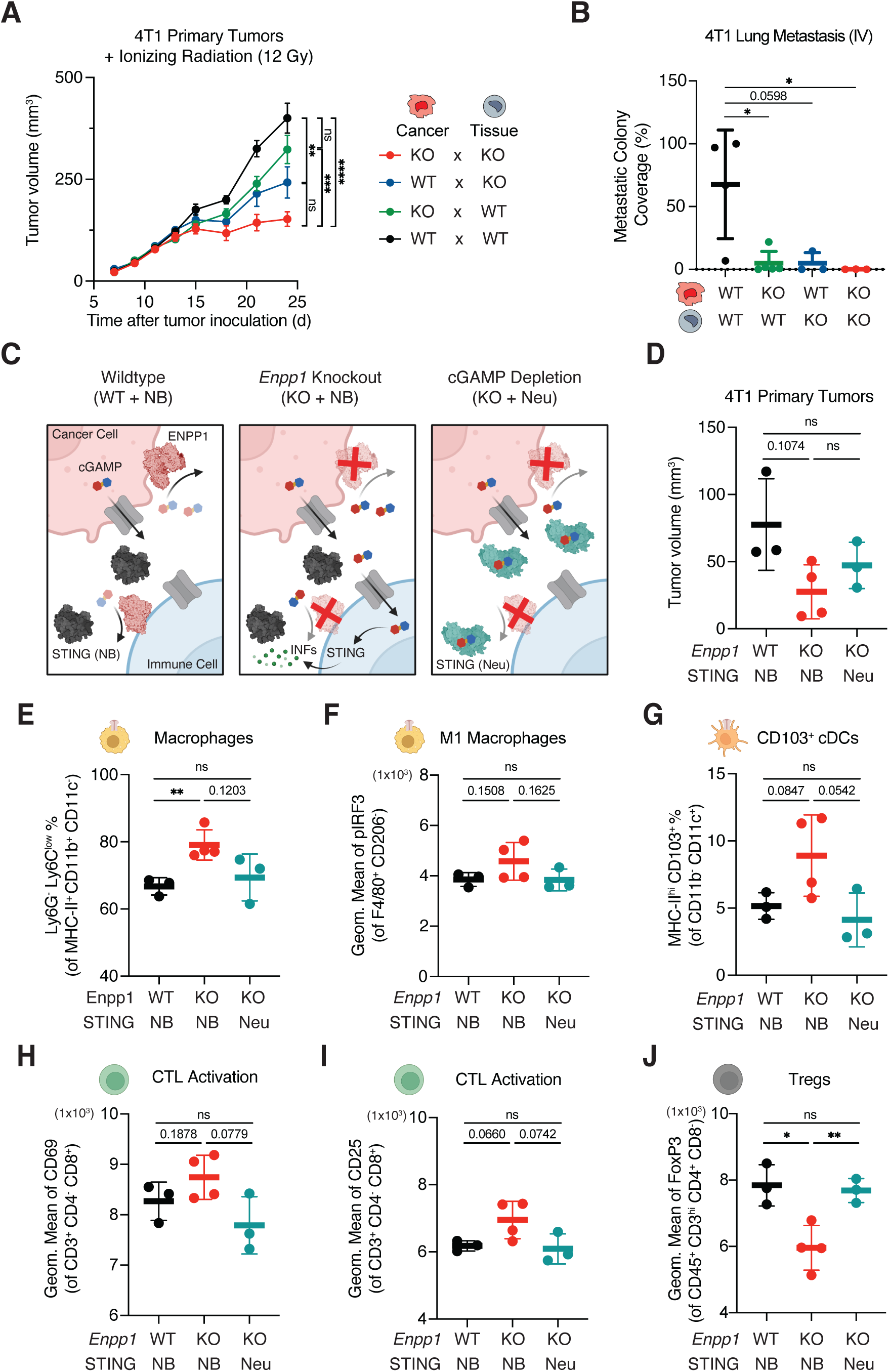
The antitumoral and immunostimulatory effect of ENPP1 deficiency is dependent on increased extracellular cGAMP. (A) Primary tumor volumes of WT or *Enpp1*^-/-^ 4T1 BALB/cJ mice orthotopically injected with WT or *Enpp1*^-/-^ 4T1 (n = 7, 17, 9, 16 mice for *Enpp1* KO x KO, KO x WT, WT x KO, and WT x WT cancer x tissue genotype combinations). Data were plotted as mean ± SEM. *P* values of the last tumor measurement were determined by multiple unpaired *t* test with Welch correction. (B) Quantifications of lung metastatic colonies of WT or *Enpp1*^-/-^ 4T1 BALB/cJ mice intravenously injected with WT or *Enpp1*^-/-^ 4T1 (n = 4, 5, 3, 3 mice for *Enpp1* KO x KO, KO x WT, WT x KO, and WT x WT cancer x tissue genotype combinations). Data were plotted as mean ± SD. *P* values were determined by unpaired *t* test. (C) Schematic of experimental strategies of comparing between wildtype, *Enpp1* knockout, and extracellular cGAMP depletion through genetic manipulation and cGAMP neutralization. Structures of dimer ENPP1 (PWD: 4B56), monomer ENPP1 (PWD: 6XKD), mSTING (PWD: 4KCO) and mSTING bound with DMXAA (PWD: 4LOL). (D) Primary tumor volumes of *Enpp1* WT mice receiving *Enpp1* WT 4T1 and R237A non-binding STING injection (WT + NB) (n = 3 biological replicates), *Enpp1* KO mice receiving *Enpp1* KO 4T1 and NB STING injection (KO + NB) (n = 4 biological replicates), and *Enpp1* KO mice receiving *Enpp1* KO 4T1 and WT neutralizing STING injection (KO + Neu) (n = 3 biological replicates). (E) The percentage of Ly6G^-^Ly6C^low^ cells out of MHC-II^+^CD11b^+^CD11c^-^ macrophages. (F) The geometric mean of pIRF3 of F40/80^+^CD206^-^ M1-like macrophages. (G) The percentage of MHC-II^hi^CD103^+^ cells out of CD11b^-^CD11c^+^ cells. (H and I) The geometric mean of CD69 (H) and CD25 (I) of CD3^+^CD4^-^CD8^+^ cytotoxic T lymphocytes (CTLs). (J) The geometric mean of Foxp3 of CD45^+^CD3^hi^CD4^+^CD8^-^ regulatory T cells (Tregs). (D-E) Data were plotted as mean ± SD. *P* values were determined by unpaired *t* test with Welch correction. \**P* ≤ 0.05., \*\**P* ≤ 0.01; *P* value is shown if between 0.05 and 0.2; not significant (ns). See also Figure S6.

We also investigated how the ENPP1 source impacts metastasis by orthotopically implanting WT or *Enpp1^-/-^* 4T1 cells into WT or *Enpp1^-/-^* 4T1 mice, respectively (WT x WT vs. KO x KO) and collected various organs for *ex vivo* culturing at experimental end point. In WT x WT mice, distal metastasis was observed in the draining inguinal lymph node (dLN), blood, lung and liver, but not in the brain. On the other hand, we observed no metastasis in KO x KO mice (**Figure S6C**). Since the KO x KO mice also have attenuated primary tumor growth, we reasoned that this could mask direct effects on metastasis, and therefore sought to disambiguate metastasis from primary tumor growth rate by intravenously injecting 4T1 cells. Again, we found that total loss of ENPP1 in cancer cells and host tissue rendered two thirds of the mice metastasis-free (**Figure 4B, S6D**). Our results support a model in which both tissue-derived and tumor-derived ENPP1 act in concert to promote primary tumor growth and distal organ metastasis. Importantly, the complete lack of detectable metastasis in most ENPP1-deficient animals harboring ENPP1-deficient tumors indicates a significant therapeutic potential of ENPP1 inhibition (Carozza, Brown, *et al*., 2020) to protect against metastasis.

### The antitumoral and immunostimulatory effect of ENPP1 deficiency Is dependent on increased extracellular cGAMP

Our scRNA-seq analysis revealed that both dampening cGAMP signaling and promoting eADO signaling contribute to an immunosuppressive primary TME in ENPP1 overexpressing tumors. When reversed to consider the ENPP1-deficient condition, we hypothesize that as less eADO will be generated, the extracellular cGAMP-STING signaling will play a dominant role in immune recruitment and activation. To test this hypothesis, we took advantage of previously developed cell-impermeable STING protein as a neutralizing agent to deplete extracellular cGAMP and compared it with R237A mutant STING that does not bind to cGAMP as a negative control (Carozza, Böhnert, *et al*., 2020; Cordova *et al*., 2021). We injected either WT neutralizing STING (Neu) or R237A non-binding STING (NB) into WT mice bearing WT 4T1 orthotopic tumors (WT x WT) or *Enpp1* KO mice with ENPP1 KO 4T1 orthotopic tumors (KO x KO) (**Figure 4C**). On day 15, we observed a near 75% reduction in tumor sizes upon *Enpp1* deletion: an effect that was around 50% rescued with extracellular cGAMP neutralization but not with the NB STING treatment (**Figure 4D)**. From these data, we conclude that *Enpp1* knockout slows primary tumor growth at least partially through enhancing extracellular cGAMP levels.

We also analyzed the immunological changes in these primary tumors in response to *Enpp1* knockout with or without extracellular cGAMP depletion by flow cytometry. We observed an increase in the percentage of macrophages in KO x KO tumors that was reversed upon extracellular cGAMP depletion (**Figure 4E**). In KO x KO tumors, we observed increased STING activation in M1-like macrophages measured by IRF3 phosphorylation (**Figure 4F**), increased number of CD103^+^ migratory cDCs (**Figure 4G**), and increased expression of activation markers CD69 and CD25 in cytotoxic T cells (**Figure 4H, I**). Additionally, we observed decreased immunosuppressive regulatory T cell (Treg) marker FoxP3, all indicating a more immunostimulatory TME (**Figure 4J**). However, these effects are completely abolished upon extracellular cGAMP depletion (**Figure 4F-J**). Together, our data provide a mechanistic link between enhanced extracellular cGAMP signaling and immunological control of primary tumor growth upon *Enpp1* loss.

### Selective inhibition of ENPP1’s cGAMP hydrolysis activity reduces breast cancer development in a STING-dependent manner

Apart from its cGAMP hydrolysis activity, ENPP1 is also known to degrade ATP (Li *et al*., 2021). While our experiments above support a model in which extracellular cGAMP signaling is at least partially responsible for the effects of ENPP1 on tumorigenesis, we wanted to formally test the sufficiency of cGAMP hydrolysis to explain ENPP1’s pro-tumorigenic phenotypes. We took advantage of the previously developed homozygous *Enpp1^H362A^* mouse model: a separation-of-function point mutant that does not degrade cGAMP but retains its catalytic activity towards ATP and other nucleotide triphosphate substrates (Carozza *et al*., 2022). Expanding beyond the 4T1 tumor model, we observed delayed primary E0771 tumor growth as measured by improved survival outcomes (time taken for tumors to reach 1000 mm^3^) in *Enpp1^H362A^* mice compared to WT mice (**Figure 5A**). *Enpp1^H362A^* mice retarded E0771 tumor growth to a similar degree as *Enpp1^-/-^* mice, as compared to WT mice (Carozza, Böhnert, *et al*., 2020), and the tumor slowing effects in *Enpp1^H362A^*mice were completely abolished in the *Sting* knockout background (**Figure 5A**).

**Figure 5.**
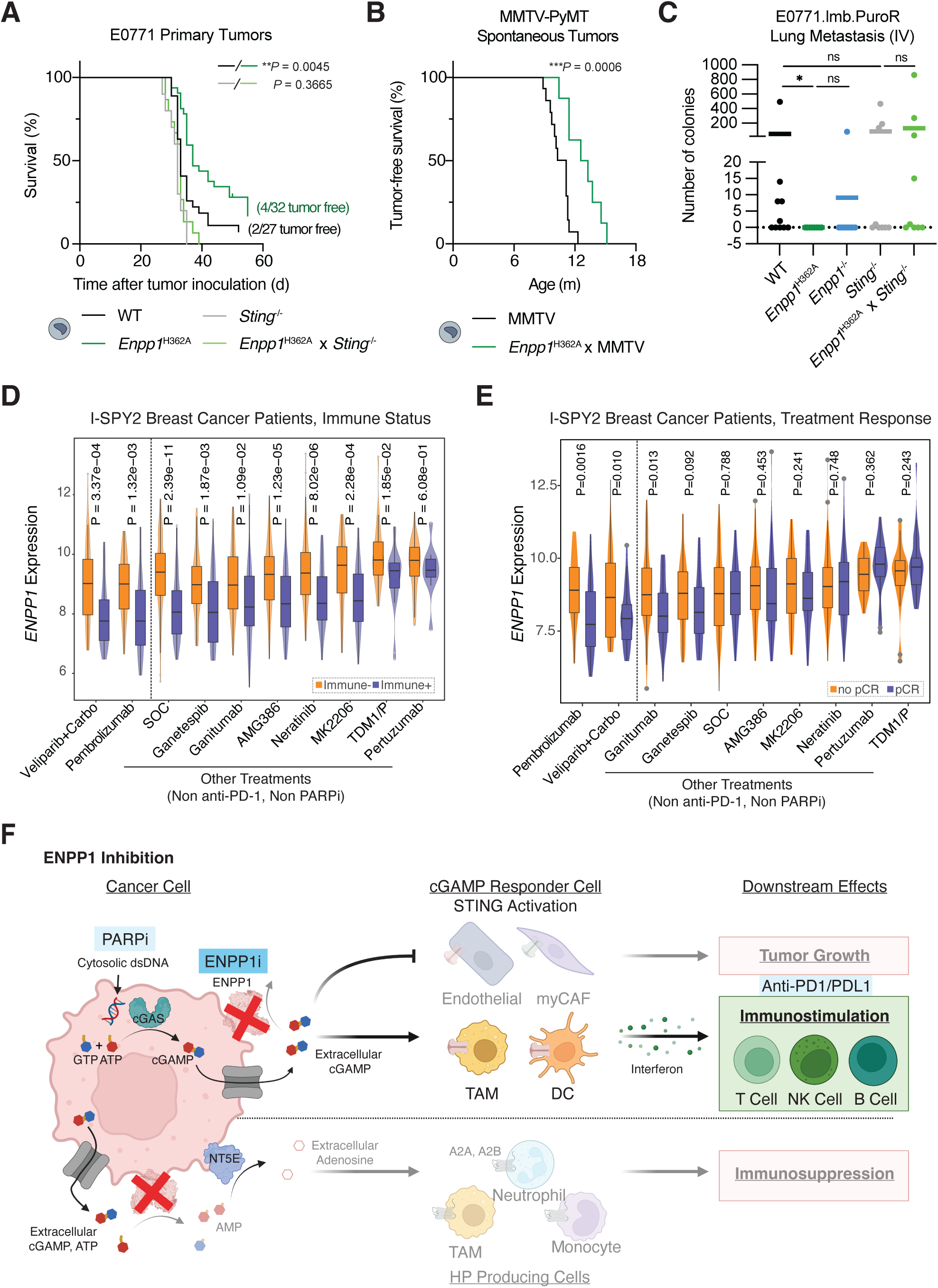
Selective inhibition of ENPP1’s cGAMP hydrolysis activity reduces breast cancer development in a STING-dependent manner. (A) Survival of WT, *Enpp1^H362A^, Sting^-/-^*, and *Enpp1^H362A^*x *Sting^-/-^* C57BL/6J mice (n = 27, 32, 10, 15 mice) bearing orthotopic E0771 breast tumors. Survival was measured by time taken for orthotopic E0771 breast tumors to reach 1000m^3^. (B) Tumor-free survival of MMTV and *Enpp1^H362A^* x MMTV mice (n = 15 and 18 mice) that developed spontaneous breast tumors. Tumor-free survival was measured by onset of the first spontaneous breast tumors. (C) Number of lung metastatic colonies of WT, *Enpp1^H362A^*, *Enpp1^-/-^. Sting^-/-^* and *Enpp1^H362A^* x *Sting^-/-^* C57BL/6J mice (*n* = 10, 8, 9, 9, 9 mice) intravenously injected with E0771.lmb.PuroR cells. Data are shown as mean. *P* value was determined by nonparametric Mann-Whitney *U* test. (D-E) *ENPP1* expression in Immune+ vs. immune-patients (D) and pCR vs. no pCR patients (E) across 10 treatment arms in the I-SPY2 breast cancer clinical trial. (F) Proposed model of mechanism of action in ENPP1 inhibition. *P* value for Kaplan-Meier curves were determined by log-rank Mantel-Cox test. \**P* ≤ 0.05, \*\**P* ≤ 0.01, \*\*\**P* ≤ 0.001. not significant (ns). Immune+ stands for immune-positive; immune-stands for immune-negative; pCR stands for pathological complete response. See also Figure S6.

Furthermore, we noticed a 69% increase in the tumor-free rate in *Enpp1^H362A^*compared with WT mice after E0771 implantation (4/32 [1.25%] vs. 2/17 [0.74%]) (**Figure 5A**). Therefore, we hypothesized that *Enpp1* depletion also disfavors primary tumor onset. To test this hypothesis, we adopted a spontaneous breast tumor model of hemizygous mice harboring the mouse mammary tumor virus-polyoma middle tumor-antigen (MMTV-PyMT, MMTV for short). The median time for female MMTV mice to develop spontaneous breast tumors is 11 months of age (**Figure 5B**). Remarkably, *Enpp1*^H362A^ x MMTV mice with impaired extracellular cGAMP degradation activity exhibited delayed median tumor onset by 2 months (**Figure 5B**). These findings support cGAMP-mediated protection from tumor initiation upon ENPP1 blockade.

Lastly, to determine whether blocking ENPP1’s cGAMP hydrolysis activity alone is sufficient for preventing metastasis, we intravenously injected E0771.lmb.PuroR (a derived metastatic cell line from the parental E0771 cells (Johnstone *et al*., 2015) engineered with puromycin resistance to allow for *ex vivo* selection) into mice of different genetic backgrounds. 50% of WT mice but none of the *Enpp1^H362A^* mice and only 11% of *Enpp1^-/-^* mice had lung metastasis. Moreover, the anti-metastatic effect of blocking ENPP1’s cGAMP hydrolysis activity in the *Enpp1^H362A^* mice is completely STING-dependent (**Figure 5C**). We postulate that deactivating ENPP1’s cGAMP hydrolysis activity to enhance paracrine STING signaling will be a promising therapeutic approach to impede breast cancer metastasis.

In addition to serving as a monotherapy, ENPP1 inhibition (ENPP1i) could potentially synergize with existing therapies that are upstream of cGAMP production, such as PARP inhibitors (PARPi) that induce DNA damage (Ding *et al*., 2018), or downstream of STING-mediated immune infiltration, such as anti-PD1/PD-L1 (Tumeh *et al*., 2014). In the I-SPY2 clinical trial (Wolf *et al*., 2022), breast cancer patients who had immune-positive (Immune+) tumors as defined by dendritic cell infiltration and STAT1 expression (downstream of IFN-I signaling) (Ivashkiv and Donlin, 2014) had significantly less *ENPP1* mRNA expression universally across all 10 treatment arms (**Figure 5D**). Strikingly, *ENPP1*-low patients gained marked pathological complete response (pCR) in Pembrolizumab (anti-PD1) and Veliparib + Carboplatin (PARPi) arms only: the only treatments that have a clear mechanistic connection to cGAMP-STING signaling. These results suggest that some patients may respond better to anti-PD1 and PARPi at least partially because of enhanced cGAMP-STING mediated immune infiltration permitted by an *ENPP1*-low setting (**Figure 5E**). Together, we put forward a model that boosting extracellular cGAMP-STING signaling is the major mechanism of action of ENPP1 inhibition in mounting immune protection against breast cancer, a strategy that could synergize with existing PARPi or anti-PD1/PDL1 therapies (**Figure 5F**).

## Discussion

Motivated by a lack of understanding of the role of paracrine cGAMP-STING signaling in cancer and contradictory mechanistic hypotheses of ENPP1’s role in cancer immunity, we aimed to uncover the causal molecular and cellular mechanisms by which ENPP1 impacts primary breast tumor growth and metastasis. While we previously identified ENPP1 as a cGAMP hydrolase, there has been significant debate as to whether its ability to deplete cGAMP and thereby dampen STING signaling is central to its pro-tumorigenic effects, as ENPP1 has other enzymatic activities toward ATP and other nucleotide triphosphates but also generates eADO – a cancer-associated metabolite – as a byproduct of its cGAMP hydrolase activity. Using an unbiased scRNA-seq approach, we systematically characterized the immunological impacts and signaling events upon overexpression of ENPP1’s catalytic activity in orthotopically implanted 4T1 cancer cells. We discovered that ENPP1-high cancer cells promote breast tumor growth by shunting the immunostimulatory cGAMP-STING pathway to the immunosuppressive eADO pathway, while fostering an angiogenic TME for tumor survival (**Figure S5D**). Using the *Enpp1^H362A^* variant that specifically abolishes ENPP1’s cGAMP hydrolysis activity and orthogonal molecular sponges to deplete extracellular cGAMP, we confirmed that cGAMP is the relevant substrate in *in vivo* cancer models in a manner dependent on downstream STING signaling. Together, our results demonstrate the importance of extracellular cGAMP-STING activation in antitumoral immunity and ascribe ENPP1 as an innate immune checkpoint of the extracellular cGAMP-STING pathway.

ENPP1’s contribution to different stages of tumor development including initiation, progression, and metastasis was not well understood. Importantly, our work provides the first evidence that ENPP1 promotes breast cancer initiation (**Figure 5B**). Comparing between primary tumors and metastases, we noticed a stronger contribution of cGAMP-STING inhibition to the pro-metastatic phenotype of ENPP1 in our scRNA-seq analyses. We posit that this could be due to elevated cGAMP production along the oncogenic trajectory, as we showed that CIN-high pro-metastatic *Kif2c*+ cancer cells (Bakhoum *et al*., 2018b) expressed higher levels of *Cgas* (**Figure 3A**). While a previous study attributed the increasing role of cGAMP hydrolysis by ENPP1 in metastasis to it replacing ATP as the major source of eADO (Li *et al*., 2021), we raise an alternative explanation that direct dampening of cGAMP-STING activation is the culprit in metastasis.

Prior studies supported a role for ENPP1 in promoting various cancer types, but largely focused on ENPP1 expressed on the cancer cells (Takahashi *et al*., 2015; Hu *et al*., 2019; Bageritz *et al*., 2014, Lau *et al*., 2013; Li *et al*., 2021). Our scRNA-seq analyses revealed that not only is ENPP1 expressed on many host responder cells, but that responder cell-derived ENPP1 has an outsized effect on blocking paracrine STING activation in those same cells. The importance of cancer- and responder cell-derived ENPP1 in tumor development is in line with our understanding of paracrine extracellular cGAMP signaling as being short-ranged. ENPP1 on the surface of cGAMP-producing and cGAMP-sensing cells would be ideally poised to snatch a freshly exported or soon-to-be imported cGAMP molecule in close proximity to cGAMP transporters (Ritchie *et al*., 2019; Lahey *et al*., 2020; Cordova *et al*., 2021), thereby circumventing paracrine activation of the STING pathway within the TME. As cGAS is rarely inactivated in cancer cells (Bakhoum *et al*., 2018b) and there is no known intracellular cGAMP hydrolase (Li *et al*., 2014), we bring forward a model that cancer cells export the high levels of cGAMP they produce, capitalizing on both cancer cell- and responder cell-derived ENPP1 for its extracellular clearance, to achieve immune evasion.

Our data highlight the notion that host-derived ENPP1 is not a passive bystander, but rather actively involved in shaping the tumor immune microenvironment. We hypothesize that the ENPP1 status of the tissue in which cancer develops, either as the primary site or the site of metastasis, together with ENPP1 allele or expression level variations, could dictate the extent of ENPP1’s role in tumor development and even the risk of tumor development altogether. Furthermore, the I-SPY2 human clinical trial data nominated ENPP1 as a potential prognostic biomarker: while ENPP1-low breast cancer patients would more likely benefit from anti-PD1 and PARPi therapies, ENPP1-high breast cancer patients may warrant combination therapy of ENPP1i with anti-PD1 or PARPi.

In summary, our work pinpoints the mechanisms and signaling events important for the anti-tumoral and anti-metastatic effects of ENPP1 depletion in several breast cancer models in mice. Future discovery of the regulatory mechanisms and genetic variants affecting ENPP1 activity, localization, and expression level in different tissues and cancer types will reveal additional strategies leading to immune evasion. As a central player dictating cancer-innate-adaptive immune communication through the STING pathway, ENPP1 is a promising target for cancer immunotherapy that may bolster our arsenal of ICB therapeutics as a druggable innate immune checkpoint.

## Methods

### Mouse strains

C57BL/6J (strain #000664), BALB/cJ (strain #000651), C57BL/6J-*Sting^gt^/J* (strain #017537), C57BL/6J-*Enpp1^asj/GrsrJ^* (strain #012810), BALB/cJ-*Enpp1^asj-2J/GrsrJ^* (strain #019107), and FVB/N-Tg(MMTV-PyVT) 634Mul/J (strain #002374) mice were purchased from the Jackson Laboratory. C57BL/6J-*Enpp1^H362A^* and C57BL/6J-*Enpp1^H362A^*x *Sting^-/-^* mice were generated and characterized in house (Carozza *et al*., 2022). FVB/N-Tg(MMTV-PyVT) 634Mul/J mice were bred with C57BL/6J-WT or C57BL/6J-*Enpp1^H362A^* to generate B6;FVB-MMTV and B6;FVB-*Enpp1^H362A^* x MMTV respectively. For MMTV spontaneous tumor model, female mice from the second generation of both B6;FVB-MMTV and B6;FVB-*Enpp1^H362A^* x MMTV genotypes were used for experiment. For all other breast tumor experiments, female mice between 6-15 weeks old were used for tumor experiments. Mice were maintained at Stanford University in compliance with the Stanford University Institutional Animal Care and Use Committee (IACUC) regulations. All procedures were approved by the Stanford University Administrative Panel on Laboratory Animal Care (APLAC).

### Mammalian cell lines and primary cells

4T1 and E0771.lmb cells were procured from ATCC. E0771 cells were procured from CH3 BioSystems. 4T1-luciferase (4T1-luc) cells were a gift from C. Contag, Stanford University, Stanford, CA, USA (Vilalta *et al*., 2014). 4T1-luc *Enpp1*^-/-^ pooled clonal cell line and 293T cGAS *ENPP1^-/-^* single clonal cell line were generated in a previous study (Carozza, Böhnert, *et al*., 2020). Primary mouse lung fibroblasts were isolated by incubating the minced lungs with 1 mg/mL collagenase from *Clostridium histolyticum* (Sigma-Aldrich) and 20 μg/mL DNase I (Sigma-Aldrich) for 1 hour, then passed through 100 μM cell strainer, spun down, treated with RBC lysis buffer for 5 minutes and plated in 10cm dish until all fibroblasts attached. To isolate metastasis from the blood or distant organs in animal studies, blood, draining inguinal lymph nodes (dLNs), lungs, livers, and brains were collected at experimental end point. Blood was dispensed into a 15-ml conical tube containing 10 mL 1x HBSS (Gibco), centrifuged at 1500 rpm at room temperature for 5 minutes before plated. dLNs were teased apart by forcing through a 100 μM strainer before plated. All other organs were minced with scissors and forces and digested in collagenase from *Clostridium histolyticum* (Sigma-Aldrich) at various conditions: 4°C for 75 minutes for lungs, 37°C for 30 minutes for livers, and 37°C for 120 minutes for brains. Digested organs were passed through a 100 μM cell strainer, washed twice with 1x HBSS (Gibco) before plating. 4T1, E0771, E0771.lmb, and their derived cell lines were maintained in RPMI (Corning Cellgro) supplemented with 10% FBS (R&D Systems), 10 mM HEPES (Gibco), and 1% penicillin-streptomycin (ThermoFisher). 293T cGAS *ENPP1^-/-^*and mouse lung fibroblast cells were maintained in DMEM (Corning Cellgro) supplemented with 10% FBS and 1% penicillin-streptomycin. 4T1 and E0771.lmb.PuroR metastasis were cultured for 7-10 days with minimal disturbance in IMDM (Gibco) supplemented with 10% FBS, 1% penicillin-streptomycin, and 60 μM 6-thioguanine (Sigma-Aldrich) or 1.5 μg/mL puromycin (Sigma-Aldrich) respectively. All cells were maintained in a humidified incubator at 37°C and 5% CO_2_. All cell lines tested negative for mycoplasma contamination.

### Synthesis and purification of cGAMP and [^32^P] cGAMP

To enzymatically synthesize cGAMP (Ritchie *et al*., 2019), 1 μM purified sscGAS was incubated testis DNA (Sigma) for 24 h. The reaction was then heated at 95°C for 3 min and filtered through a 3-kDa filter. cGAMP was purified from the reaction mixture using a PLRP-S polymeric reversed phase preparatory column (100 Å, 8 μm, 300 x 25 mm; Agilent Technologies) on a preparatory HPLC (1260 Infinity LC system; Agilent Technologies) connected to UV-vis detector (ProStar; Agilent Technologies) and fraction collector (440-LC; Agilent Technologies). The flow rate was set to 25 mL/min. The mobile phase consisted of 10 mM triethylammonium acetate in water and acetonitrile. The mobile phase started as 2% acetonitrile for the first 5 min. Acetonitrile was then ramped up to 30% from 5-20 min, then to 90% from 20-22 min, maintained at 90% from 22-25 min, and then ramped down to 2% from 25-28 min. Fractions containing cGAMP were lyophilized and resuspended in water. The concentration was determined by measuring absorbance at 280 nm. To enzymatically synthesize [^32^P] cGAMP, 1 μM purified sscGAS was incubated with 20 mM Tris-HCl pH 7.4, 250 μCi (3000 Ci/mmol) [α-^32^P] ATP (Perkin Elmer), 1 mM GTP, 20 mM MgCl_2_, and 100 μg/mL herring testis DNA (Sigma) in a reaction volume of 100 μL for 24 h. The reaction was purified by preparatory TLC on a HP-TLC silica gel plate (Millipore), eluted in water, and filtered through a 3-kDa filter to remove silica gel.

### STING expression and purification

WT neutralizing STING or R237A non-binding STING were expressed and purified using previously published methods (Carozza, Böhnert, *et al*., 2020). In brief, pTB146 His-SUMO-mSTING (residues 139-378) was expressed in Rosetta (DE3) pLysS competent cells (Sigma-Aldrich). Cells were grown in 2xYT medium with 100 μg/mL ampicillin until they reached an OD600 of 1. They were then induced with 0.75 mM IPTG at 16°C overnight. Cells were pelleted and resuspended in 50 mM Tris pH 7.5, 400 mM NaCl, 10 mM imidazole, 2 mM DTT, and cOmplete protease inhibitors (Sigma-Aldrich). The cells were then flash frozen and thawed twice before sonication to lyse the cells. The supernatant was incubated with HisPur cobalt resin (Thermo Scientific) for 30 min at 4°C. The resin-bound protein was washed with 50 column volumes of 50 mM Tris pH 7.5, 150 mM NaCl, 2% triton X-114; 50 column volumes of 50 mM Tris pH 7.5, 1 M NaCl; and 20 column volumes of 50 mM Tris pH 7.5, 150 mM NaCl. Protein was eluted from resin with 600 mM imidazole in 50 mM Tris pH 7.5, 159 mM NaCl. Fractions containing His-SUMO-STING were pooled, concentrated, and dialyzed against 50 mM Tris pH 7.5, 150 mM NaCl while incubating with SUMOlase His-ULP1 to remove the His-SUMO tag overnight. The solution was incubated with the HisPur cobalt resin again to remove the His-SUMO tag, and STING was collected from the flowthrough. Protein was dialyzed against 20 mM Tris pH 7.5, loaded onto a HitrapQ anion exchange column (GE Healthcare) using Äkta FPLC (GE Healthcare), and eluted with a NaCl gradient. Fractions containing STING were pooled, buffer exchanged into PBS, and stored at −80°C until use.

### Recombinant DNA

To clone guide RNA plasmid targeting *Enpp1*, *Enpp1* guide sequences listed in **Table S1** were cloned into the BbsI site of PX458-GFP or P458-mCherry backbones (synthesized by Addgene) following the protocol from Ryuji Morizane Lab at Harvard in order. To clone pLenti-CMV-mENPP1-WT-GFP-Puro plasmid, mENPP1-WT sequence was amplified from pcDNA3-mENPP1-FLAG (synthesized by Genscript) using pLenti_mENPP1_fwd and pLenti_mENPP1_rev primers in **Table S1** and inserted into the XbaI-BamHI sites of pLenti-CMV-GFP-Puro (Addgene). To clone pLenti-CMV-mENPP1-T238A-GFP-Puro plasmid, T238A point mutants were first introduced into pcDNA3-mENPP1-FLAG using QuikChange mutagenesis. mENPP1-T238A sequence was then introduced into the XbaI-BamHI sites of pLenti-CMV-GFP-Puro. pLenti-TetONE-FLAG-Puro were synthesized by Addgene and used to generate E0771.lmb.PuroR cell line. All oligonucleotide sequences are in **Table S1**.

### Generation of transiently edited cell lines

4T1 *Enpp1^-/-^* cells were created the same way as 4T1-luc *Enpp1^-/-^* cells previously described (Carozza, Böhnert, *et al*., 2020). Briefly, 4T1 underwent transient transfection with Lipofectamine 3000 of the following pairs of sgRNAs targeting mouse *Enpp1*: PX458-mENPP1_sgRNA1-GFP and PX458-mENPP1_sgRNA2-mCherry; PX458-mENPP1_sgRNA3-GFP and PX458-mENPP1_sgRNA1-mCherry. Double GFP and mCherry positive cells were FACS sorted and underwent single-cell cloning. Sequence knockout was confirmed with PCR using mENPP1_sgRNA12_seq_fwd and mENPP1_sgRNA12_seq_rev, or mENPP1_sgRNA34_seq_fwd and mENPP1_sgRNA34_seq_rev primer pairs. Functional knockout was confirmed with activity assay (commercial antibodies are not sensitive enough for verification of protein expression). Multiple clean knockout clones were pooled to generate the 4T1 *Enpp1^-/-^* cell line.

### Generation of stable expression cell lines

4T1 *Enpp1^-/-^* cells generated above were then virally transfected to stably express WT or T238A mouse ENPP1, giving rise to ENPP1^WT-OE^ or ENPP1^T238A-OE^ cell lines. Briefly, lentiviral packaging plasmids (pHDM-G, pHDM-Hgmp2, pHDM-tat1b, and pRC.CMV-rev1b) were purchased from Harvard Medical School. 500 ng of pLenti-CMV-mENPP1-GFP-Puro, pLenti-CMV-mENPP1-T238A-GFP-Puro, or pLenti-CMV-mENPP1-A84S-GFP-Puro plasmid, and 500 ng of each of the packaging plasmids were transfected into 293T cells using FuGENE 6 transfection reagent (Promega). The viral media was exchanged after 24 h, harvested after 48 h and passed through a 0.45 μm filter, and used to transduce 4T1 *Enpp1^-/-^* cells (4T1 cells are used as 4T1-luc cells already carry puromycin resistance which will interfere with subsequent drug selection process). 48 hours later, cells were selected with 1–2 μg/ml puromycin, single-cell cloned, and 4-6 clones were pooled after verification by western blot and activity assay. E0771.lmb cells were virally transduced with empty pLenti-TetONE-FLAG-Puro vector to stably carry puromycin resistance following the previously described lentiviral transfection protocol, giving rise to the E0771.LMB.PuroR cell line used in metastatic models.

### Serum and lysate preparation

Mouse blood was collected through terminal cardiac puncture and spun at 2,000 x g for 5 min. The resulting serum layer was collected and stored at −80°C until use. Cell lysate preparation for cGAMP degradation activity assay is as following: cells from a confluent well in a 6 well plate was collected in 1 mL of PBS, centrifuged at 1000 x *g* for 3 minutes, lysed in 50-100 μL of lysis buffer (10 mM Tris pH 9, 150 mM NaCl, 10 μM ZnCl_2_, 1% NP-40), and stored in −20°C until use. For western blotting, cells were lysed on the plate in 100-250 μL of Laemmli sample buffer, boiled at 95°C for 5 minutes and sonicated. Mouse tumor lysate (100 mg/mL) for cGAMP degradation activity assay were generated by lysing tissues in 10 mM Tris pH 7.5, 150 mM NaCl, 10 μM ZnCl_2_, 1.5% NP-40, and freshly added protease inhibitors (cOmplete, EDTA-free protease inhibitor cocktail, Sigma-Aldrich). Organ lysates were then homogenized with a bead homogenizer (Omni International) and stored at −20°C until use.

### [^32^P] cGAMP degradation thin-layer chromatography assays

For experiments reported in **Figure S1A-C**, cGAMP degradation activity assay (20 μl) for cells containing 50% cell lysate, cGAMP (1 μM, with trace [^32^P] cGAMP spiked in), and standard ENPP1 activity buffer (50 mM Tris pH 9, 250 mM NaCl, 0.5 mM CaCl_2_, 1 μM ZnCl_2_) took place in room temperature. For experiments reported in **Figure S6A-B**, cGAMP degradation activity assay for moues organs (30 μl) containing 75% organ lysate (100 mg/mL), 5 μM cGAMP, and PBS took place in 37°C. Lastly, cGAMP degradation activity assay (20 μl) for mouse serum containing 50% serum, 5 μM cGAMP and physiological ENPP1 activity buffer (50 mM Tris pH 7.5, 150 mM NaCl, 1.5 mM CaCl_2_, 10 μM ZnCl_2_) took place in 37°C. At indicated times, 1 μl aliquots of the reaction were quenched by spotting on HP-TLC silica gel plates (Millipore). The TLC plates were run in mobile phase (85% ethanol, 5 mM NH_4_HCO_3_) and exposed to a phosphor screen (GE BAS-IP MS). Screens were imaged on a Typhoon 9400 scanner and the ^32^P signal was quantified using ImageJ. The sample size and statistical tests of computation are indicated in respective figure legend.

### Western blotting

Cell lysates were separated on an SDS-polyacrylamide gel (Genscript) and transferred to a nitrocellulose membrane using a wet transfer system (BioRad). Primary antibody mouse anti-tubulin (1:2000) and rabbit anti-GFP (1:1000) were purchased from Cell Signaling and added overnight at 4°C, followed by three washes in TBS-T (1x TBS-0.1% tween). Secondary antibody IRDye 800CW goat anti-rabbit (1:15,000) and IRDye 680RD goat anti-mouse (1:15,000) were purchased from Li-COR Biosciences and added for 1 h at room temperature, followed by three additional washes in TBS-T. Blots were imaged in IR using a LI-COR Odyssey Blot Imager. Bands were quantified using ImageJ.

### 4T1 murine breast tumor models

For experiment reported in **Figure 1**, BALB/cJ female mice were orthotopically injected with 2.5×10^6^ ENPP1^WT-OE^ or ENPP1^T238A-OE^ 4T1 suspended in 100 μL of PBS cells in the 4^th^ mammary fat pad (MFP). When tumors reached 1000 mm^3^, we sacrificed the animals and collected primary tumors and lungs. Tissues were processed into single cell suspension following steps described above. Half of the lung suspension were plated into 60 μM 6-thioguanine (Sigma-Aldrich) and 10% heat-inactivated FBS (R&D Systems) containing IMDM (ThermoFisher) media and cultured for 6-12 days without disturbance. At the end of the experiment, colonies of metastases were fixed in methanol and visualized with 0.03% (w/v) methylene blue (Sigma-Aldrich). Metastatic colonies were quantified with Fuji Image J. The rest half of the lung suspension and the tumor suspension were cryopreserved and later thawed for scRNA-seq. For primary tumor experiment reported in **Figure 4A** and **Figure S6A-B**, we orthotopically injected 5 x 10^4^ WT or *Enpp1^-/-^* 4T1-luc cells into WT or *Enpp1^-/-^* BALB/cJ. When tumors were palpable with an average tumor volume of 100 ± 20 mm^3^ (determined by length^2^ x width / 2), 10-12 days after cell inoculation, tumors were irradiated with 12 Gy using a 225 kVp cabinet X-ray irradiator filtered with 0.5 mm Cu (IC-250, Kimtron Inc., CT) following previously described procedures (Carozza, Böhnert, *et al*., 2020). We sacrificed mice when their tumors reached 1000 mm^3^ and collected their primary tumors and sera for cGAMP degradation activity following steps described above. For tumor metastasis experiment reported in **Figure 4B, S4D**, we intravenously injected 5 x 10^4^ WT or *Enpp1^-/-^* 4T1 cells into the tail veins of WT or *Enpp1^-/-^*BALB/cJ mice. We collected their lungs around day 30 and quantified metastatic burden following process describe above. For tumor metastasis experiment reported in **Figure S4C**, we orthotopically injected 2.5 x 10^4^ WT or *Enpp1^-/-^* 4T1 cells into the tail veins of WT or *Enpp1^-/-^* BALB/cJ mice, respectively. At day 33, we sacrificed the animals and collected blood, draining inguinal lymph nodes, primary tumors, lungs, livers, and brains for metastasis assay. The sample size and statistical tests of computation are indicated in respective figure legend.

### E0771 murine breast tumor models

For primary tumor experiment in **Figure 5A**, we orthotopically injected 2.5 x 10^4^ E0771 cells into the 4^th^ MFP of WT, *Enpp1^H362A^*, *Sting^-/-^,* or *Enpp1^H362A^* x *Sting^-/-^* C57BL/6J mice. We measured animal survival by the time it took for the tumors to reach 1000 mm^3^. For tumor metastasis experiment in **Figure 5C**, we intravenously injected E0771.lmb.PuroR cells into the tail veins of WT, *Enpp1^H362A^*, *Enpp1^-/-^*, *Sting^-/-^,* and *Enpp1^H362A^*x *Sting^-/-^* C57BL/6J mice. 30 days later, we cultured dissociated lungs in 1μg/μL puromycin for 9 days before methylene blue visualization. The sample size and statistical tests of computation are indicated in respective figure legend.

### MMTV spontaneous breast tumor model

Female mice hemizygous for the MMTV-PyMT transgene and homozygous for the wild type or H362A mutated *Enpp1* gene from the second generation were included in experiments. Individual tumor volume (determined by 0.5 x length x width^2^) was added up to yield total tumor volume. Tumor onset was defined by the date on which a palpable tumor of more than 0.5 mm^3^ was observed. Tumor-free survival was plotted, and statistical significance was assessed by log-rank Mantel-Cox test.

### FACS analysis of tumors upon *Enpp1* knockout and/or cGAMP depletion

5 x 10^4^ WT or *Enpp1^-/-^*4T1-luc cells were orthotopically injected into WT or *Enpp1 ^-/-^*BALB/cJ mice respectively. Starting the next day, mice were intratumorally injected with 100 μL of 100 μM neutralizing (WT) or non-binding (R237A) STING every other day up to day 13. Mice were euthanized on day 15 and tumors were collected and digested in RPMI + 10% FBS with 20 μg/ml DNase I type IV (Sigma-Aldrich) and 1 mg/ml Collagenase from Clostridium histolyticum (Sigma-Aldrich) at 37 °C for 30 min. Tumors were passed through a 100 μm cell strainer (Fisher Scientific) and red blood cells were lysed using red blood cell lysis buffer (155 mM NH_4_Cl, 12 mM NaHCO_3_, 0.1 mM EDTA) for 5 min at room temperature. Cells were stained with Live/Dead fixable nearIR or blue dead cell staining kit (ThermoFisher). Samples were then fixed and permeabilized with eBioscience Foxp3/Transcription Factor Staining Buffer Set (Invitrogen), Fc-blocked for 10 min using TruStain fcX (BioLegend) and subsequently antibody-stained with antibodies. Cells were analyzed using a Symphony (BD Biosciences), or an Aurora analyzer (Cytek). Data was analyzed using FlowJo V10 software (BD) and Prism 9.1.0 software (Graphpad) for statistical analysis and statistical significance was assessed using the unpaired *t* test with Welch’s correction.

### ScRNA-seq of murine primary tumors and lungs with metastases

Mammary fat pad primary tumors and lungs containing metastases were harvested from two ENPP1^WT-OE^ or ENPP1^T238A-OE^ tumor bearing mice when primary tumors reached 1000 mm^3^ (**Figure 1**). Cryopreserved single cell suspensions were thawed into warm RPMI media with 10% HI-FBS. Cells were washed with PBS and 0.04% w/v BSA, passed through 40 μm Flowmi cell strainer (Bel-Art, 974-25244), and assessed for concentration and viability with an automated cell counter. Viability varied between 72-87% for tumor samples and 91-96% for lung samples. Cells were resuspended in PBS and 0.04% w/v BSA to 1000 cells/μL. Single-cell suspensions were processed with the Chromium Controller microfluidic device (10X Genomics), using the Chromium Next GEM Single Cell 3’ HT Reagent Kits v3.1 (Dual Index). Library was performed according to the manufacture’s instructions (single cell 3’ HT reagent kits v3.1 protocol, 10x Genomics). Briefly, cells were resuspended in the master mix and loaded together with partitioning oil and gel beads into the chip to generate the gel bead-in-emulsion (GEM). The poly-A RNA from the cell lysate contained in every single GEM was retrotranscripted to cDNA, which contains an Ilumina R1 primer sequence, Unique Molecular Identifier (UMI) and the 10x Barcode. The pooled barcoded cDNA was then cleaned up with Silane DynaBeads, amplified by PCR and the appropriately sized fragments were selected with SPRIselect reagent for subsequent library construction. During the library construction Ilumina R2 primer sequence, paired-end constructs with P5 and P7 sequences and a sample index were added. Pooled libraries were sequenced on NextSeq 2000 (Illumina) targeting 20,000 reads per cell. Sequencing data was demultiplexed, mapped to the murine reference genome (Ensembl release 93 GRCm38), and gene counts were quantified per cell using Cell Ranger (version 3.1.0.). In total, we obtained 81,694,726 reads across 4,830 cells, and detected on average (median) 1,445 genes per cell.

### ScRNA-seq clustering and cell type annotation

All subsequent processing, quality control, and analyses were performed with Cellenics (https://scp.biomage.net/data-management). We filtered out cells with high mitochondrial content and high doublet count. We then applied Harmony data integration with HVGs = 2000 and performed dimensionality reduction using 27 principal components (90.18% variation). We performed Louvain clustering with resolution set to be 0.8. We manually annotated 18 clusters based on known cell markers (**Figure S2A**). Specifically, we identified cancer cells based on markers enriched in 4T1 compared to BALB/cJ mammary fat pad (Schrörs *et al*., 2020). A separate cancer cluster was annotated for overexpressing *Kif2c*, which has been reported to drive CIN and metastasis (Bakhoum *et al*., 2018a) and additional genes related to metastasis (Schrörs *et al*., 2020). Among fibroblasts, a unique cluster enriched in tumors overexpresses myofibroblast marker *Acta2*, as well as *Tgfb1* and *Tgfb2* and are known as myofibroblastic cancer-associated fibroblasts (myCAF) and overtly immunosuppressive (Mhaidly and Mechta-Grigoriou, 2020; Mao *et al*., 2021). We also performed detailed subcluster analysis on macrophages (**Figure S3A-B**) and T cells (**Figure S3C-D**). Cluster frequency analysis, gene expression analysis, and differential gene expression are performed in Cellenics.

### ISPY-2 patient dataset analysis

Normalized, batch corrected microarray data and sample metadata were retrieved from GSE194040. Individual two-tailed t-tests were performed comparing ENPP1 expression between patients who achieved a pathological complete response (pCR) and those who did not, within each ISPY-2 clinical arm. Similarly, ENPP1 expression was compared between tumors that were likely to respond to immunotherapy (immune+) and those that were not using two-tailed t-tests. Immune+status was estimated based on the average dendritic cell and STAT1 signatures (Wolf *et al*., 2022).

### Quantification and statistical analysis

In cGAMP degradation assays, half-life was obtained by one phase exponential decay fitting with Prism software, with intercept (Y0) set up to be the initial cGAMP concentration, and plateau set to be 0. In proliferation assays, proliferation rate was obtained by exponential fitting with Prism software. All statistical tests were performed using GraphPad Prism software and are noted in the figure legends. Data are presented as the mean ± standard deviation unless otherwise stated.

## Lead contact

Further information and requests for resources and reagents should be directed to and will be fulfilled by the lead contact, Lingyin Li

## Materials availability

The plasmids and the ENPP1^WT-OE^ and ENPP1^T238A-OE^ 4T1 cell lines will be made available upon request.

## Data and code availability

- scRNA-seq raw and processed data can be accessed via GSE233659
- All data reported in this paper will be shared by the lead contact upon request.
- Any additional information required to reanalyze the data reported in this paper is available from the lead contact upon request.

## Acknowledgement

We thank J. Carozza for providing primary mouse lung derived fibroblasts, and for generating 4T1 *Enpp1*^-/-^ and 4T1-luc *Enpp1*^-/-^ pooled clonal cell lines, as well as 293T cGAS *ENPP1*^-/-^ clonal cells lines. We thank A. Pawluk for constructive feedback on the manuscript. We thank all Li Lab members for their constructive comments and discussion through the course of this study. S.W. was supported by the Stanford Medical Scholars Fellowship, Stanford Berg Scholars Program, and Stanford Chi-Li Pao Foundation Alpha Omega Alpha Student Research Fellowship. A.J.J. was supported by NSF Graduate Research Fellowship Program. V.S. was supported by Stanford Graduate Fellowship. H.G. is an Era of Hope Scholar (W81XWH-2210121). L.A.G. is supported by a NIH New Innovator Award (DP2 CA239597), a Pew-Steward Scholars for Cancer Research award, and the Arc Institute. This work was supported by NIH DP2CA228044 (L.L)., NIH 5R01CA258427-02 (L.L.)., and Arc Institute.

## Author contributions

S.W. conceived the project, performed the experiments, analyzed results, coordinated the project, and wrote the research manuscript. V.B. conceived the project and performed the experiments. A.J.J. and V.S. supported with interpreting results, performing experiments, and analyzing results. J.S., F.B.Y., and L.A.G. supported with the scRNA-seq experiment and analyses. X.L. supported with performing animal experiments. G.S. helped bred mice used for the experiments. V.S and H.G. performed METABRIC and I-SPY2 human data analyses. H.G. and L.A.G. both provided scientific expertise and approved the final research manuscript. L.L., as leading senior author, conceived the project, coordinated the project, and provided the project funding. S.W., A.J.J., V.S., H.G. and L.A.G. all helped with funding acquisition.

