## Supplemental Information for "ENPP1 is an innate immune checkpoint of the anticancer cGAMP-STING pathway"

**Supplemental Table 1. Oligonucleotide Sequences**

| Name | Sequence (5'->3') |
|---|---|
| **Primers for cloning** | |
| mENPP1_T238A_fwd | GCCTATGTACCCTACCAAGgcgTTTCCCAATCATTACAGC |
| mENPP1_T238A_rev | GCTGTAATGATTGGGAAAcgcCTTGGTAGGGTACATAGGC |
| mENPP1_seq_fwd | CTACAGTTCTGTGTGCCAAG |
| mENPP1_seq_rev | CCATTATTGGGAGCTGGGATCAAACC |
| pLenti_mENPP1_fwd | gccatccacgctgttttgacctccatagaagacaccgactctagaGATCCGCCACCATGGAGC |
| pLenti_mENPP1_rev | aacagctcctcgcccttgctcaccatggtggcgaccggtggatccCAGAATTCGTCTTCTTGGCTGAAGATTG |
| **For CRISPR editing cells** | |
| mENPP1_sgRNA1_for | caccgGCTCGCGCCCATGGACCT |
| mENPP1_sgRNA1_rev | aaacAGGTCCATGGGCGCGAGCc |
| mENPP1_sgRNA2_for | caccgATATGACTGTACCCTACGGG |
| mENPP1_sgRNA2_rev | aaacCCCGTAGGGTACAGTCATATc |
| mENPP1_sgRNA3_for | caccgGTGACCTGCAGGGTCCTT |
| mENPP1_sgRNA3_rev | aaacAAGGACCCTGCAGGTCACC |
| mENPP1_sgRNA4_for | caccgTGTGTAGCTGTGACACTCAG |
| mENPP1_sgRNA4_rev | aaacCTGAGTGTCACAGCTACACA |
| mENPP1_sgRNA12_seq_fwd | GCCAAATACCCGGGGCGTTG |
| mENPP1_sgRNA12_seq_rev | AAATCGGCGTCCTGTTTCCAGAAG |
| mENPP1_sgRNA34_seq_fwd | GGGTAGTGTGACAATAATTTATGG |
| mENPP1_sgRNA34_seq_rev | CAGGAGGGGGATATAAATGCC |

#### Supplemental Figure Legends

**Figure S1. Generation of ENPP1$^{WT\text{-}OE}$ and ENPP1$^{T239A\text{-}OE}$ 4T1s and optimization of *ex vivo* metastasis culture**

(A) ENPP1 degradation activity in 4T1 and its derived cell lines as assessed by TLC.
(B) ENPP1-GFP expression (top) and ENPP1 degradation activity (bottom) of ENPP1$^{WT\text{-}OE}$ 4T1 and ENPP1$^{T238A\text{-}OE}$ 4T1 clones as assessed by western blotting and TLC. Bolded clones were pooled for experiments.
(C) ENPP1-GFP degradation activity (left) and ENPP1 expression (right) of ENPP1$^{WT\text{-}OE}$ 4T1 and ENPP1$^{T238A\text{-}OE}$ 4T1 pooled clones as assessed by TLC and western blotting.
(D) Proliferation of ENPP1$^{WT\text{-}OE}$ and ENPP1T$^{238A\text{-}OE}$ 4T1 pooled clonal cell lines compared with WT 4T1 cells over time (n = 3 biological replicates).
(E) Experimental schematic of *ex vivo* culture of 4T1 metastasis by orthotopic injection.
(F) Number of live 4T1 cells (left) or mouse lung fibroblasts (right) 6 and 9 days after 0, 30, or 60 $\mu$M 6-thioguanine (6-TG) treatment.
Data were plotted as mean $\pm$ SD. *P* values were determined by unpaired *t* test.
TLC stands for thin layer chromatography.

**Figure S2. ScRNA-seq analysis of 4T1 murine primary tumors and lung metastases**

(A) Bubble heatmap showing expression of selected marker genes for cluster annotation. Dot size indicates fraction of expressing cells, colored based on average expression levels.
(B) UMAP plot of the annotated clusters of ENPP1$^{T238A\text{-}OE}$ and ENPP1$^{WT\text{-}OE}$ 4T1 primary tumors and metastasis colonized lungs.
(C) Barplots comparing non-immune cell compositions (containing C01-C05) between ENPP1$^{T238A\text{-}OE}$ and ENPP1$^{WT\text{-}OE}$ 4T1 primary tumors and metastasis colonized lungs.
(D and E) Violin plots of indicated transcripts of indicated cell types comparing between ENPP1$^{T238A\text{-}OE}$ and ENPP1$^{WT\text{-}OE}$ 4T1 tumors or metastases. *Arg1* in monocytes (C09) in primary tumors and lung metastases (D); *Itgae* and *H2-Ab1* in cDC1s (C12) in lung metastases (E). *P* values were determined by nonparametric Mann-Whitney *U* test.
cDC1 stands for conventional dendritic cell type 1.

**Figure S3. Subclustering of macrophages and T cells**

(A) UMAP plot of the annotated subclusters of macrophages (C10).
(B) Bubble heatmap showing expression of marker genes for macrophage subcluster annotation and functional markers indicating M1-like versus M2-like macrophage cell phenotypes. Dot size indicates fraction of expressing cells, colored based on average expression levels.
(C) Barplots comparing macrophage subcluster compositions (containing Ma1-Ma4) between ENPP1$^{T238A\text{-}OE}$ and ENPP1$^{WT\text{-}OE}$ 4T1 primary tumors and metastasis colonized lungs.
(D) UMAP plot of the annotated subclusters of T cells (C14).
(E) Bubble heatmap showing expression of marker genes for T cell subcluster annotation. Dot size indicates fraction of expressing cells, colored based on average expression levels.

(F) Barplots comparing T cell subcluster compositions (containing T01-T11) between ENPP1$^{T238A-OE}$ and ENPP1$^{WT-OE}$ 4T1 primary tumors and metastasis colonized lungs. *P* values were determined by unpaired *t* test. $*P \leq 0.05$.; *P* value is shown if it is between 0.05 - 0.15.

**Figure S4. Expression of additional genes in the STING pathway**
(A) Violin plots of *Irf3*, *Tbk1*, *Infra1*, *Ifitm1*, and *Ifitm3* across the annotated clusters.
(B-C) Bar graphs of indicated transcripts in indicated cell types comparing between ENPP1$^{T238A-OE}$ and ENPP1$^{WT-OE}$ 4T1 tumors or metastases. *Ifitms* and *Vegfc* in endothelial cells (C04) in lung metastases (B); *Lrrc8c* in macrophages (C10) in primary tumors and lung metastases.
Bars represent mean ± SEM. *P* values were determined by nonparametric Mann-Whitney *U* test.

**Figure S5. Contribution of eADO pathway and HP secretion in ENPP1 overexpression**
(A) Violin plots of *Entpd1*, *Nt5e*, *Adora2a* and *Adora2b* across the annotated clusters.
(B-C) Bar graphs of indicated transcripts in indicated cell types comparing between ENPP1$^{T238A-OE}$ and ENPP1$^{WT-OE}$ 4T1 tumors or metastases. *Adora2b*, *Tgfb1*, *Il10rb*, *Hp* in macrophages (C10) in primary tumors and lung metastases (B); *Hp* in cancer cells (C01), Kif2c+ cancer cells (C02), neutrophils (C08), and monocytes (C09) in primary tumors and lung metastases (C).
(D) Proposed model of mechanism of action in ENPP1 overexpression.
Bars represent mean ± SEM. *P* values were determined by nonparametric Mann-Whitney *U* test.
HP stands for Haptoglobin. AM stands for tumor-associated macrophages. cDC stands for conventional dendritic cell. myCAF stands for myofibroblastic cancer-associated fibroblast.

**Figure S6. *Enpp1* knockout in cancer and tissue cells**
(A-B) Representative TLC image of *ex vivo* cGAMP hydrolysis by tumor lysates (A) and relative cGAMP hydrolysis activity calculated from kinetic analysis by tumor lysates and sera (B) from randomly selected mice in Figure 4A reaching experimental endpoint (n = 2-6 biological replicates). Data represent mean ± SD. *P* values were determined by unpaired *t* test with Welch correction.
(C) Images of metastatic colonies of WT or *Enpp1*$^{-/-}$ 4T1 BALB/cJ orthotopically injected with WT or *Enpp1*$^{-/-}$ 4T1 respectively.
(D) Representative images of lung metastatic colonies and the percentage of mice with lung metastasis from Figure 3B. WT or *Enpp1*$^{-/-}$ 4T1 BALB/cJ mice were intravenously injected with WT or *Enpp1*$^{-/-}$ 4T1 (n = 4, 5, 3, 3 mice for *Enpp1* KO x KO, KO x WT, WT x KO, and WT x WT cancer x tissue genotype combinations).
(E) Representative images of lung metastatic colonies and the percentage of mice with lung metastasis from Figure 6C. WT, *Enpp1*$^{H362A}$, *Enpp1*$^{-/-}$. *Sting*$^{-/-}$ and *Enpp1*$^{H362A}$ x *Sting*$^{-/-}$ C57BL/6J mice (*n* = 10, 8, 9, 9, 9 mice) were intravenously injected with E0771.lmb cells through the tail veins. P values were determined by chi-squared test.
TLC stands for thin layer chromatography.

### FIgure S1. Generation of ENPP1$^{WT-OE}$ and ENPP1$^{T238A-OE}$ 4T1s and optimization of *ex vivo* metastasis culture

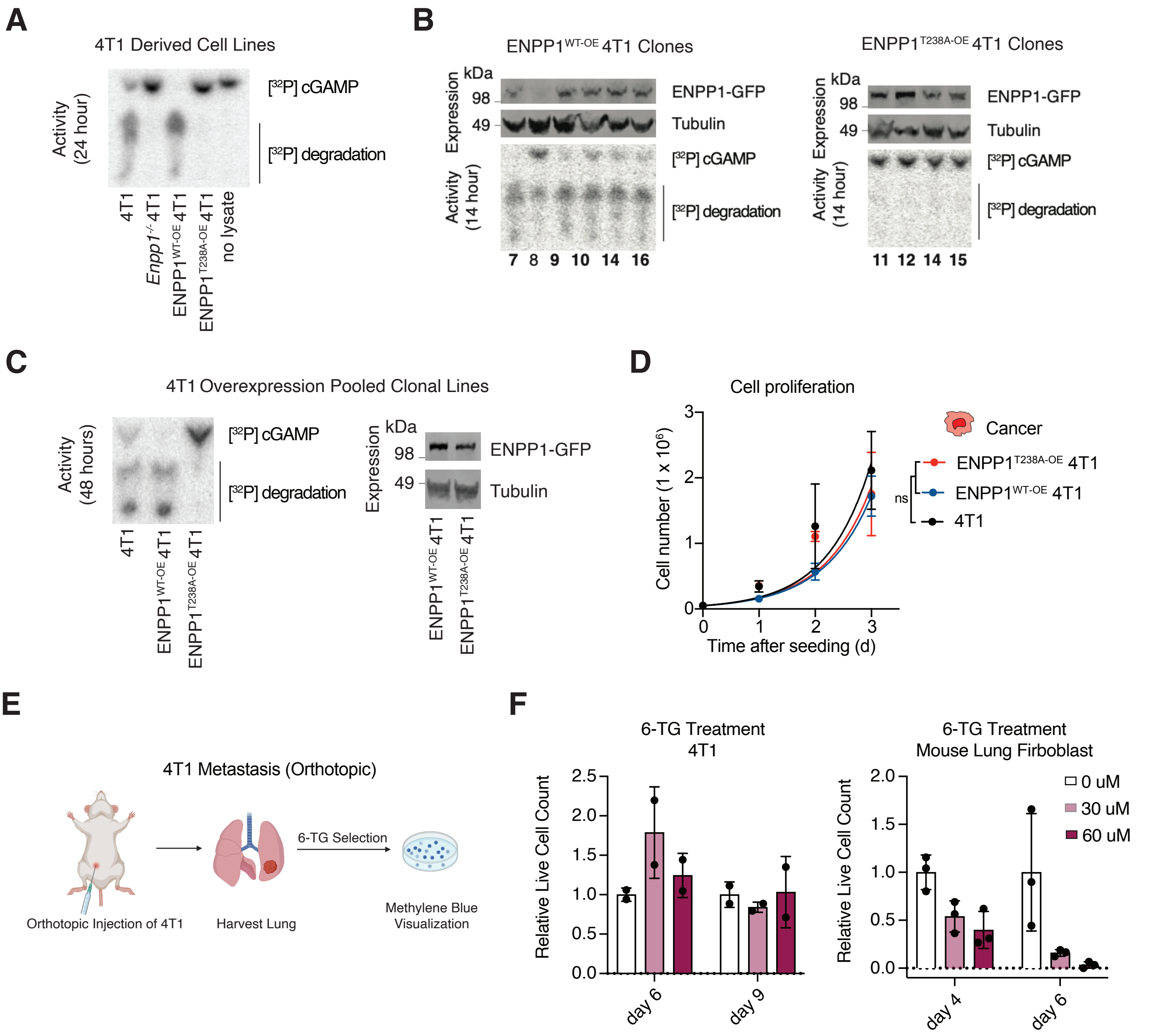

### FIgure S2. scRNA-seq analysis of murine 4T1 primary tumors and lung metastases

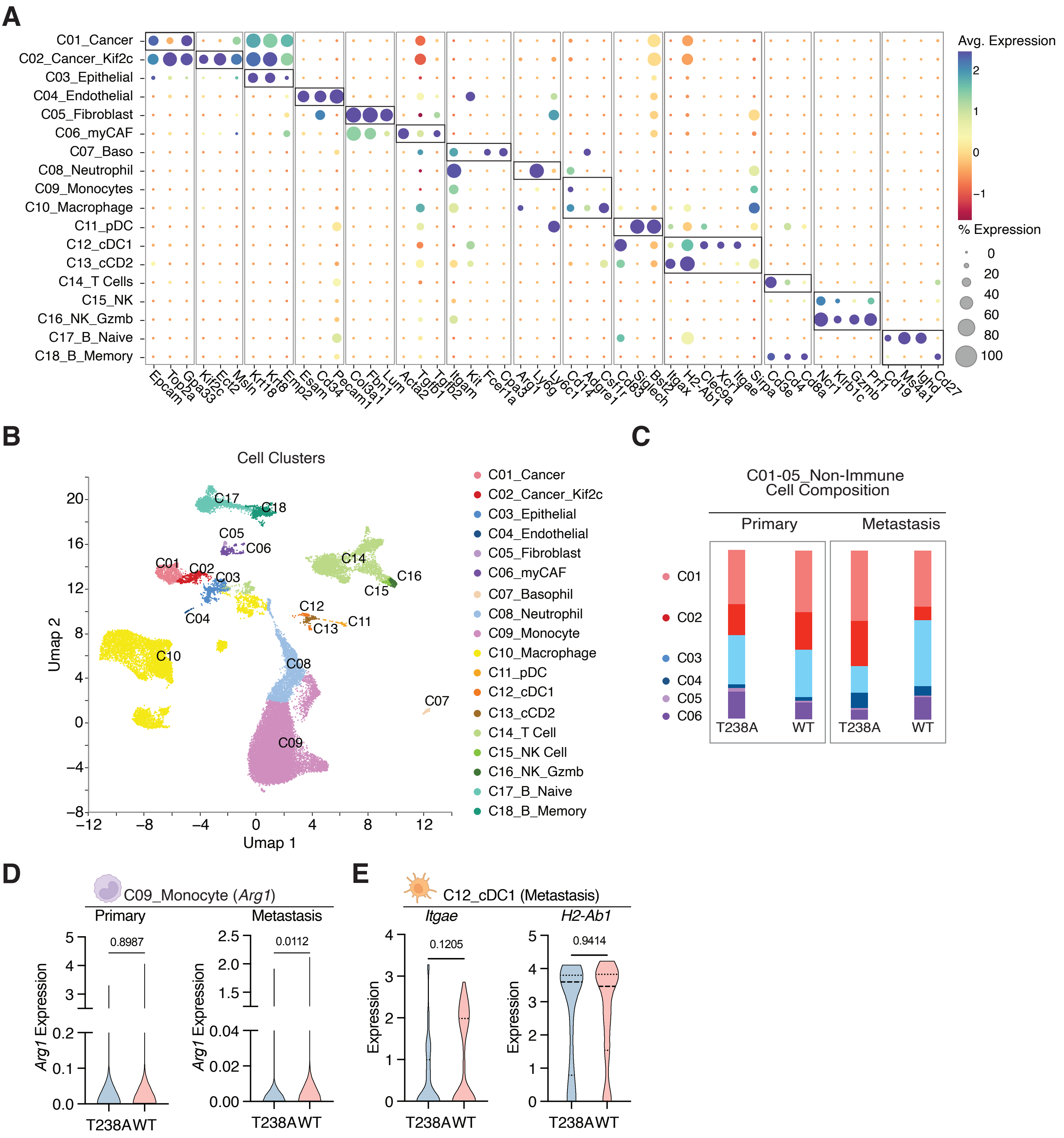

### FIgure S3. Subclustering of macrophages and T cells

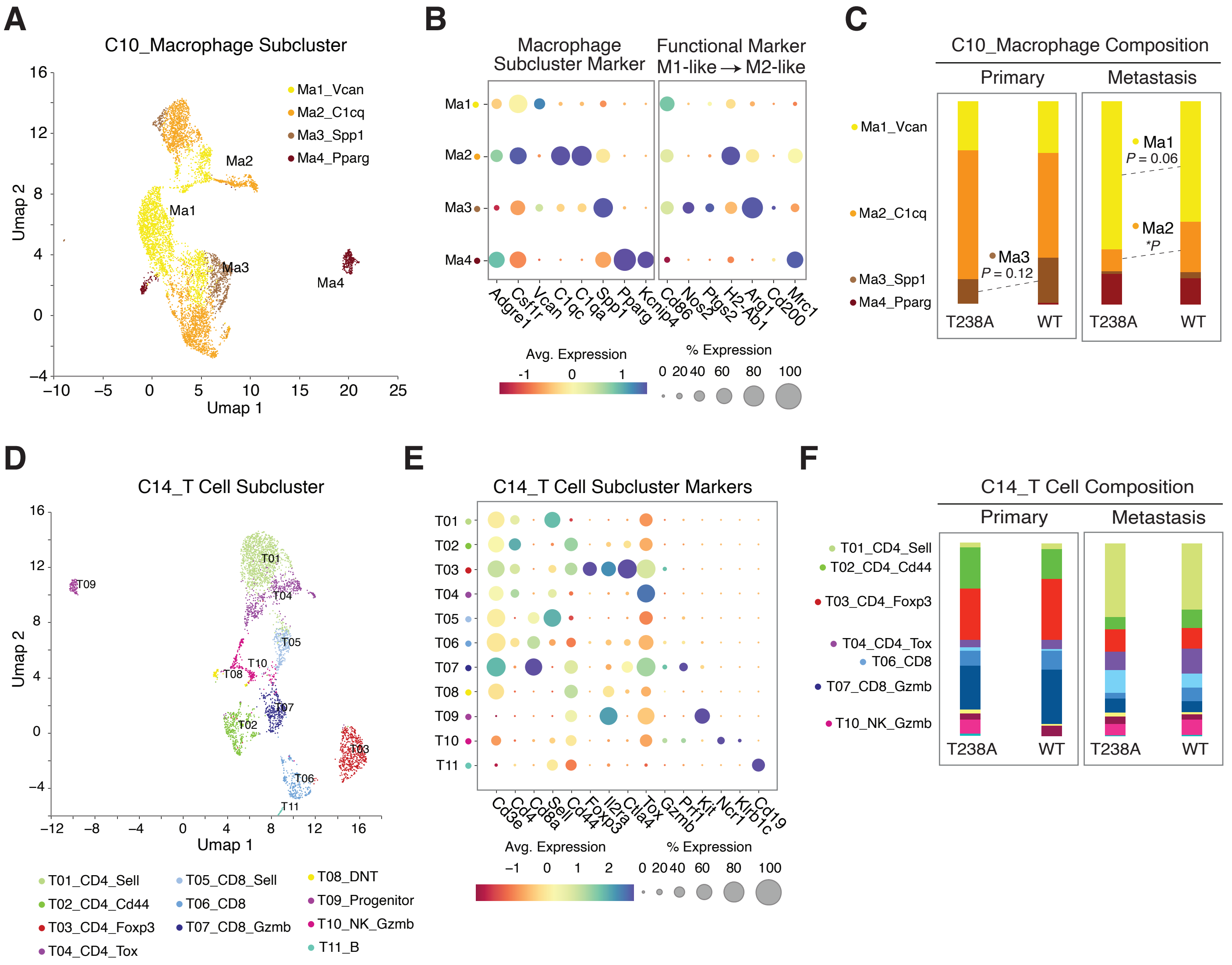

### FIgure S4. Expression of genes in the STING pathway

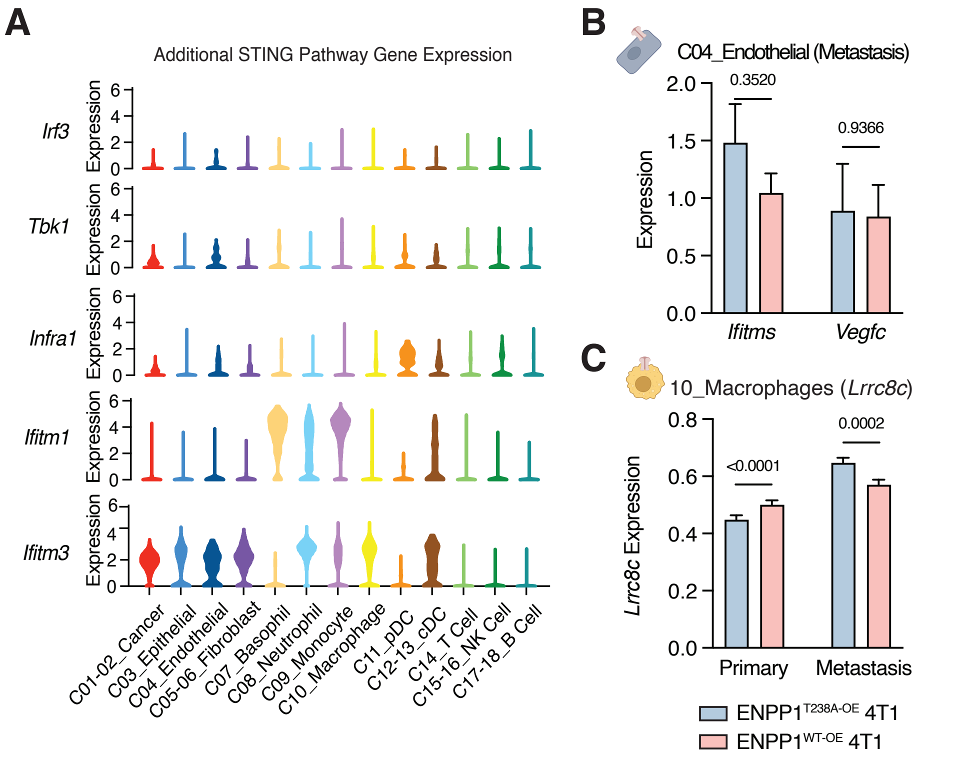

### FIgure S5. Contribution of eADO pathway and HP secretion in ENPP1 overexpression

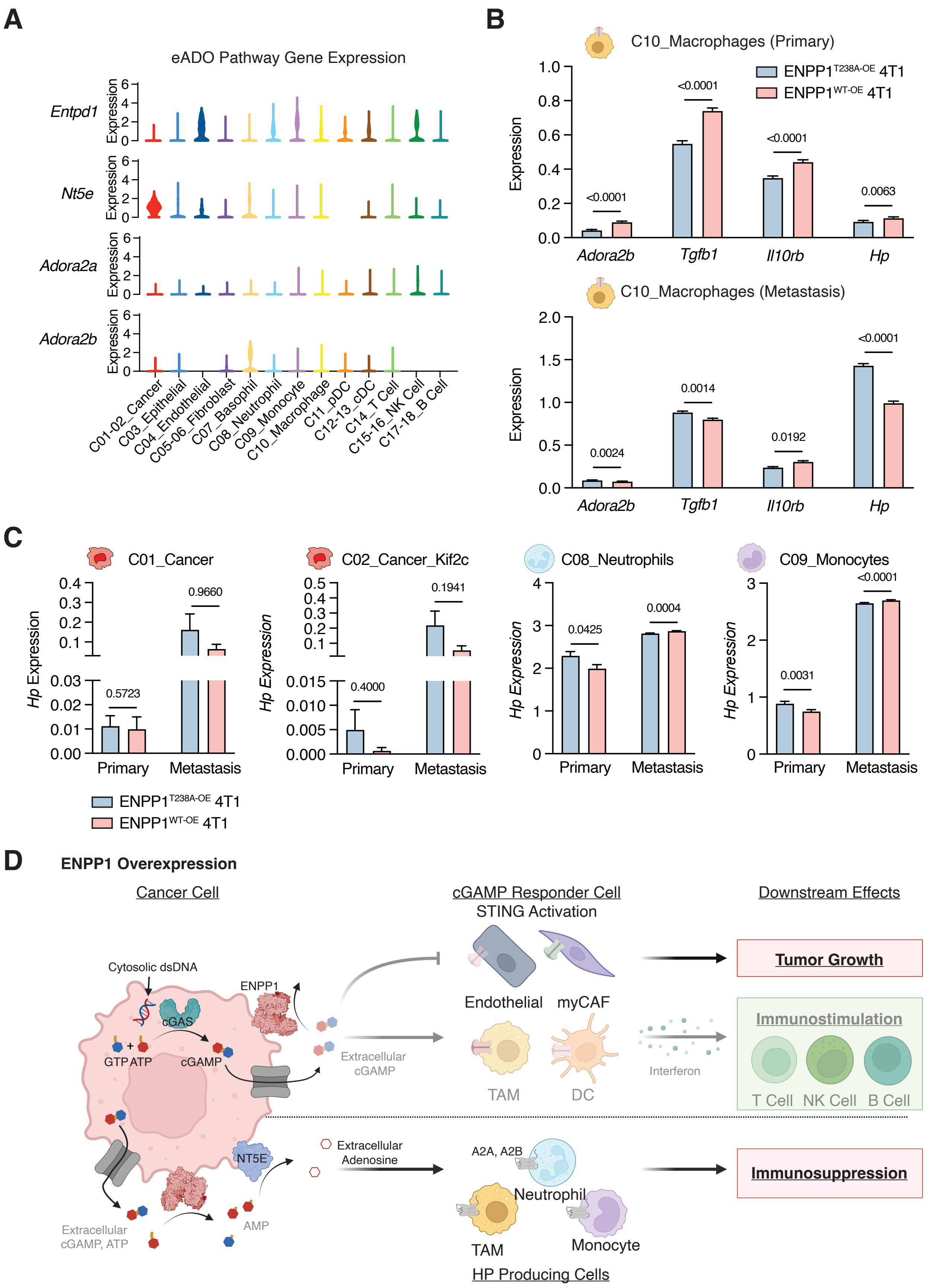

### Figure S6. *Enpp1* knockout in cancer and tissue cells

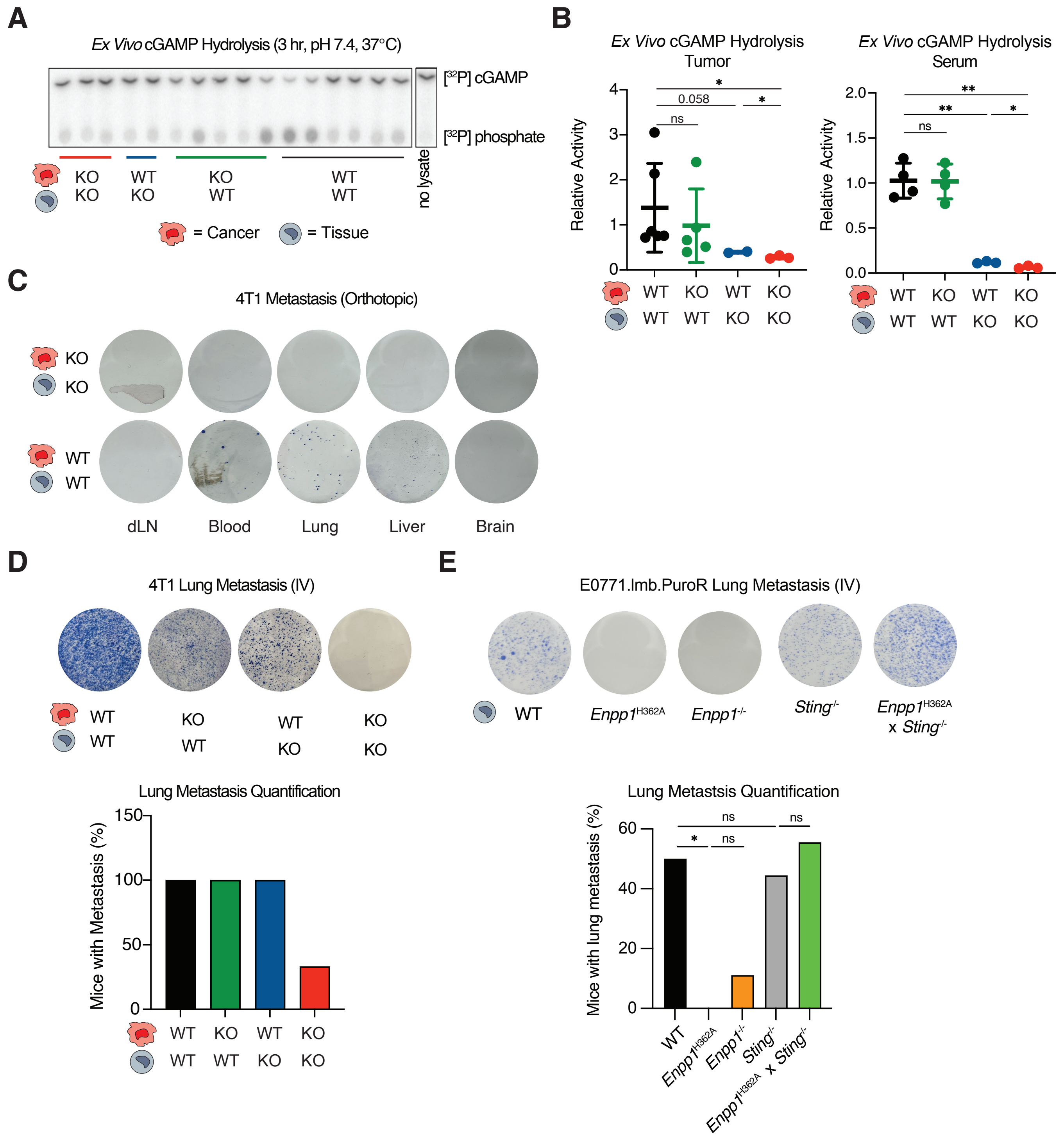
